# A single transcriptional regulator is crucial for the adaptation of *Staphylococcus aureus* to diverse niches

**DOI:** 10.1101/2025.08.01.668060

**Authors:** Claus Vogl, Lynette C. Mikula, Helene Marbach, Katharina Mayer-Weihrauch, Sebastian Herndler, Leonie Breitschopf, Amy C Pickering, Mary-Elizabeth Jobson, Adriana Cabal Rosel, Nora Dinhopl, Igor Loncaric, Werner Rupitsch, Lindsey N Shaw, Orla Keane, Ross Fitzgerald, Monika Ehling-Schulz, Simon Heilbronner, Tom Grunert

## Abstract

The adaptation of versatile multi-host pathogens to various hosts and various niches within hosts is often still poorly understood. The alternative Sigma factor B (SigB) is the master regulator of the general stress response of most gram-positive bacteria, which is a classic case of adaptive plasticity. In *Staphylococcus aureus*, SigB appears co-opted to function as a switch between intracellular and extracellular niches. During bovine mastitis, low SigB-activity confers an advantage in the milk-rich extracellular niche of the bovine udder. We show that narrowly adapted SigB-deficient strains evolved repeatedly from phenotypically plastic SigB-wildtype strains during persistent mastitis. This genetic assimilation appears driven by the cost of phenotypic plasticity: long time lags in adapting to milk and slow growth. Surprisingly, we observe that mutations causing SigB-deficiency allow even human isolates to grow in milk. While host adaptation often proceeds by mobile genetic elements exchanged between strains, we show how a master regulator in the core genome can drive niche adaptation.

## Introduction

The gram-positive bacterium *Staphylococcus aureus* is a commensal and a clinically important pathogen that can adapt to various hosts and infect various organs within these hosts. This remarkable adaptability is partly due to its large genetic variability. Strains can be differentiated according to their core genome and assigned to clonal complexes (CCs), in spite of variability in their accessory genome. CCs are often associated with a specific host. When CCs are associated with different hosts the accessory genome is host-specific and mediated by the exchange of mobile genetic elements (*1–3*). In addition to its large genetic variability, *S. aureus’s* adaptive plasticity allows it to survive and replicate in both intra- and extracellular niches (*4–6*). Sigma factor B (SigB), a member of the sigma factor family of transcriptional regulators, is crucial to this adaptability.

In *S. aureus*, the *sigB* operon contains the genes *rsbU, rsbV, rsbW,* and *sigB*. Regulation of SigB activity is complex: RsbW antagonises SigB, RsbV antagonises RsbW, and RsbU activates RsbV. Thus, the levels of *sigB* mRNA or SigB protein do not necessarily predict SigB activity; instead, transcription of SigB regulated genes is typically taken as a proxy for SigB activity (*7–10*). High SigB activity is known to induce traits necessary for intracellular survival and growth of *S. aureus*, e.g., the production of adhesins, which facilitate invasion of host cells, and of carotenoid pigments, which detoxify the reactive oxygen species within host cells (*5, 6, 11*). In contrast, low SigB activity induces secretion of virulence factors and proteases (*12, 13*), traits necessary for extracellular survival and growth. SigB thus seems to act as a switch between extra- and intracellular niches in the derived pathogen *S. aureus*. We note that SigB orchestrates a general stress response in ancestral saprophytic relatives, e.g., *Bacillus subtilis* (*14*), while *S. aureus* lacks homologs to the stressosome proteins RsbR, RsbS, and RsbT and to the energy stress-sensing proteins RsbP and RsbQ of *Bacillus subtilis* (*14, 15*). In *S. aureus*, the transcriptional regulator SigB thus seems to have been co-opted from its ancestral function in the regulation of the stress response to a derived function as switch between extra- and intracellular niches.

The two most common globally distributed bovine-associated CCs are CC97 and CC151 (*16*): CC97 strains typically cause only mild, subclinical intramammary infection (IMI) and generally have high adhesion and internalisation capacities, indicative of adaptation to the intracellular niche (*17–19*). CC151 strains cause more severe inflammation and udder damage, likely due to the secretion of extracellular virulence factors (*20, 21*). We postulate that quantitative differences in SigB activity between strains of CC151 and CC97 cause these niche-specific adaptations.

The extracellular niche in the bovine udder can be considered a chemostat: continuous secretion of milk replenishes the nutrients essential for bacterial growth, and milking removes the bacteria and their metabolites. Overall, this creates a relatively stable environment in which the increased secretion of proteinases induced by low SigB activity could benefit *S. aureus* growth and long-term survival. Persistent IMI by *S. aureus* in dairy cows may last for the host’s lifetime, allowing for short-term bacterial evolution. In fact, we have repeatedly isolated SigB-deficient (SigB-def) strains from cows with IMI and have once observed the evolution of a SigB-def strain during persistent IMI (*22–24*). We thus propose that SigB-def strains are better adapted to growth in milk. Conversely, passaging a SigB-wt strain repeatedly in the intracellular niche of cultivated macrophages produced a hyperpigmented phenotype that resulted from a mutation that canalised (i.e., fixed) constitutive overexpression of SigB (SigB-co) (*25*). SigB-def and SigB-co strains thus seem narrowly adapted to the extracellular or intracellular niche, respectively, while phenotypically plastic SigB-wt strains seem able to switch between these niches. Furthermore, while human isolates likely have never encountered the milk-rich extracellular niche of the bovine udder during their evolution, SigB-def mutations may allow even them to grow in milk.

In this article, we therefore investigate the role of SigB in the adaptation of *S. aureus* to the milk-rich extracellular niche in the bovine udder. We specifically characterise and compare the adaptation of naturally isolated (bovine and human associated) and engineered phenotypically plastic SigB-wt and canalised SigB-def strains to milk-rich media. We show that adaptive plasticity and genetic regulation of SigB activity governs a large part of *S. aureus* niche adaptation.

## Results

### General experimental set-up that enables measuring adaptation to milk

Since the focus of this article is on an ecological shift between complex niches, our experiments involve the transfer of *S. aureus* strains from one complex medium to another, specifically from tryptic soy broth (TSB) to semi-skimmed milk broth (SSM). Generally, *S. aureus* strains pre-cultivated in TSB and then transferred to fresh TSB immediately continue to grow, while the same strains transferred from TSB to SSM show an adaptation phase (i.e., a period of slow growth) until they are fully adapted to the new medium and their growth accelerates (Fig. 1A). We therefore define two target variables, similar to those proposed for monitoring bacterial growth in minimal media (*26, 27*), to assess growth after transfer from one complex medium to another: (i) the duration of the post-transfer adaptation phase (“adaptation time” symbolised by τ), which is the initial, possibly multiphasic, slow-growing phase immediately after the strains are moved from TSB to SSM, and (ii) the maximum (absolute) growth rate (symbolised by *ρ*_max_) that is reached once the strain is able to optimally utilise the mix of components in the new medium and has therefore (phenotypically) adapted to it (Fig. 1B and Material and Methods).

**Fig. 1.**
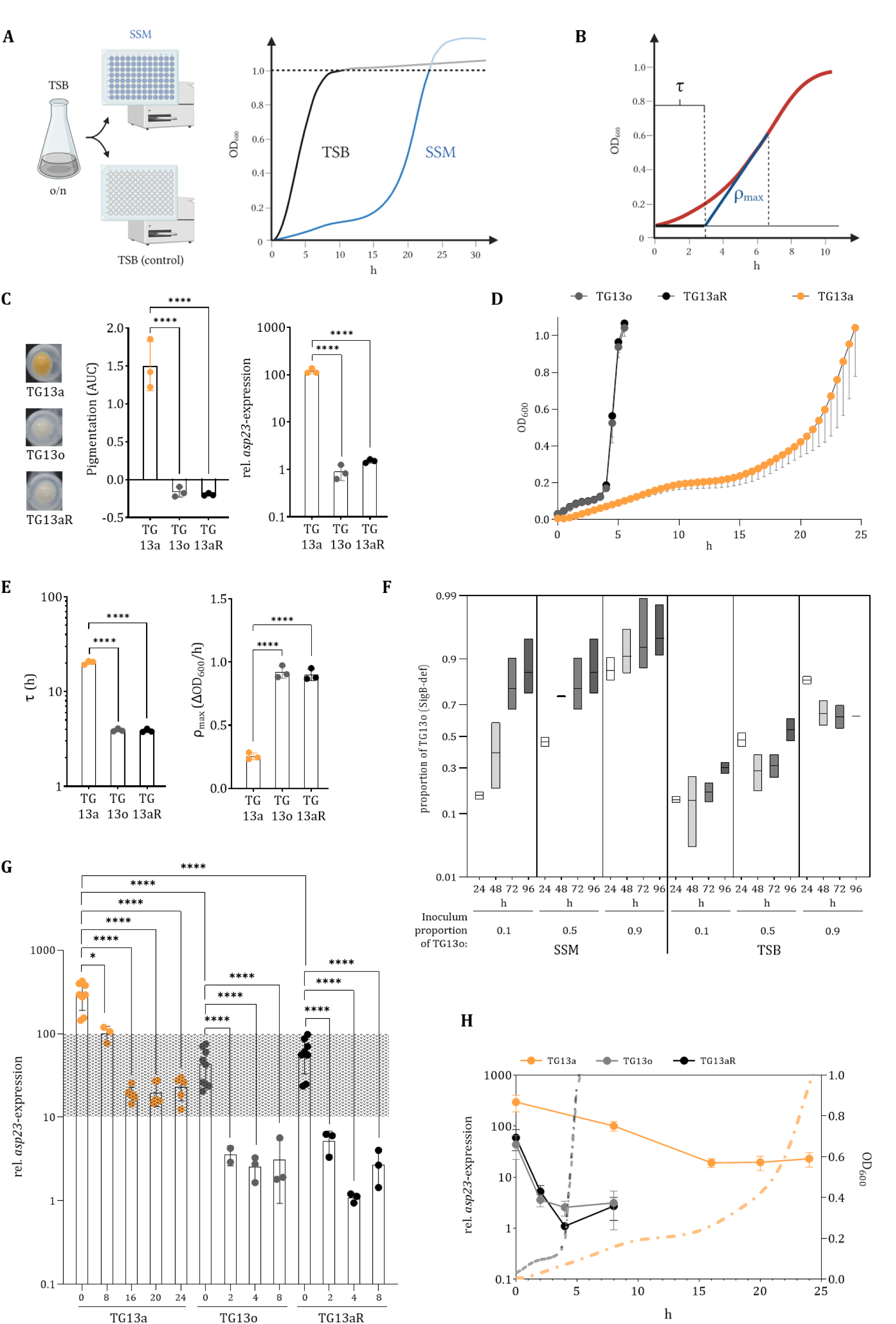
A SigB-deficient isolate grows better in milk. **(A)** Monitoring of bacterial growth in complex media up to 72 h: After transfer to fresh TSB (black line), *S. aureus* grows immediately; after transfer to SSM (blue line), it undergoes an adaptation phase before maximal growth sets in. Since milk clotting may interfere with measurement of the OD and the relationship between the OD and bacterial density deviates from linearity with increasing OD_600_, we excluded readings with OD > 1.0 from our analyses. **(B)** Calculation of the parameters “adaptation time” and “maximum growth rate” from OD_600_ time series data: From the empirical curve (red), differences of OD_600_ between neighbouring time points are calculated; the maximum difference corresponds to the maximum growth rate *ρ*_max_ (in units of ΔOD_600_/h). The line thus defined is extended to the intersection with the x-axis. The length of time from the start of the experiment to this intersection is taken as the adaptation time *τ* (in units of h). **(C – E)** Comparisons of TG13a (SigB-wt) and TG13o and TG13aR (both SigB-def) via linear model analyses of untransformed (pigmentation; *τ*) or log-transformed (RT-qPCR, *ρ*_max_) data. Differences in pairwise means were tested with Tukey’s HSD**. (C)** Proxies of *S. aureus* SigB activity: (left) Carotenoid pigmentation of the bacterial pellet at the bottom of the tube after overnight culture and centrifugation, (centre) measured by absorbance (AUC; mean ± SD), and (right) relative *asp23* mRNA expression (RT-qPCR; mean ± SD) after transfer to fresh TSB and harvest at OD_600_ = 3. **(D)** Growth dynamics in SSM after transfer from TSB measured by optical density at a wavelength of 600nm (OD_600_; mean ‒ SD). **(E)** Adaptation time (*τ*; mean ± SD) and maximum growth rate (*ρ*_max_; mean ± SD) after transfer to SSM. **(F)** Boxplots of the proportion of the derived SigB-deficient strain TG13o after 24, 48, 72 and 86 h in SSM and TSB, depending on the proportion of TG13o (vs TG13a) in the initial inoculum. **(G)** Relative *asp23* mRNA expression (RT-qPCR; mean ± SD) in the interval (h) after transfer from TSB to SSM. Differences in pairwise means were tested with Tukey’s HSD. The overlapping range of SigB activity in SigB-wt and SigB-def strains is underlaid. **(H)** Simultaneous representation of the changes in relative *asp23* mRNA expression (RT-qPCR; mean ± SD) and bacterial growth after transfer to SSM (OD_600_; mean ± SD). Relative *asp23*-expression is shown during the adaptation time (TG13a: ≅ 20h; TG13o and TG13aR: ≅ 4 h). TSB, tryptic soy broth; SSM, semi-skimmed milk broth. Fig. 1 A and B were created in https://BioRender.com.

### A SigB-def strain that evolved in the mammary gland of its host is better adapted to growth in milk than its SigB-wt ancestor

With this experimental set-up and analysis approach, we first monitor the growth of SigB-wt and SigB-def *S. aureus* strains collected from a single naturally infected dairy cow during persistent IMI. The SigB-def strain TG13o (CC97) evolved from the original SigB-wt strain TG13a by a point mutation in *rsbU*(G368A) (*22*), and obviously outcompeted the original isogenic strain over time. The same SigB-def mutation was introduced into the SigB-wt TG13a to produce the control strain TG13aR (*24*) (Supplementary Table S1). When grown in TSB, the carotenoid pigmentation of TG13o and TG13aR (both SigB-def) is low and very similar; that of TG13a (SigB-wt) considerably higher. The same pattern is reflected by the expression levels of the *asp23* gene, which is a well-established proxy of SigB activity (*7, 8, 24*) (Fig. 1C). Overall, these results indicate that the single nucleotide change in the *rsbU* gene causes SigB deficiency. When the strains are pre-cultivated in TSB and transferred to SSM, the adaptation time τ is much longer for the SigB-wt than the SigB-def strains (Fig. 1D, E and for TSB (control) see Supplementary Fig. S1A). Additionally, the SigB-def strains show a higher maximum growth rate *ρ*_max_ than the SigB-wt TG13a (Fig. 1E). Solid curd forms at the bottom of the sample vials of the SigB-def strains (TG13o and TG13aR) after 5.5 ± 1.2 h (mean ± SD) but only after 19.4 ± 1.3 h in vials of the SigB-wt TG13a strain. Counting colony-forming units (CFU/ml) confirms that the observed differences in τ and *ρ*_max_ reflect the actual growth pattern and are not driven by clotting or curd formation (Supplementary Fig. S1B). Moreover *in vitro* competition experiments confirm that the *in vivo* evolved SigB-def strain (TG13o) outcompetes the original SigB-wt strain (TG13a) in SSM alone, but not in TSB. For all ratios of inoculum, the proportion of the SigB-def strain TG13o increases over time in SSM, while changes over time are inconsistent in TSB. This is reflected by a non-significant overall regression of the proportion of the SigB-def strain on time in TSB (but a significant interaction with the initial inoculum proportion, *p* = 0.047), while the interaction between the regressors for time and the SSM medium is significantly different from zero (*p* = 0.0006) (Supplementary Table S2). In other words, the SigB-def strain outcompetes the SigB-wt strain after transfer from TSB to SSM irrespective of the initial proportion of the two strains, while neither strain has a consistent advantage in TSB (Fig. 1F).

Next, we assess the regulatory adaptation of SigB by monitoring *asp23* expression of the strains during the adaptation and rapid growth phases. Initially, expression of *asp23* is higher in the SigB-wt strain compared to the SigB-def strains (Fig. 1G and H). After transfer to SSM, the SigB-def strains show a rapid decrease in *asp23* expression to a low level that is maintained thereafter, while the SigB-wt strain shows a gradual decrease to a relatively high level. The lowest level of SigB expression is reached just before or at the time of maximum growth by all strains, however the absolute times differ: about 8 h for the SigB-def strains (TG13o and TG13aR) and about 24 h for the SigB-wt strain (TG13a) (Fig. 1H). We note that while under identical conditions *asp23* expression is lower in the SigB-def strains (TG13o and TG13aR) than in the SigB-wt strain, however, absolute levels may overlap (Fig. 1G) when comparing different conditions between strains.

Overall, the SigB-def mutation causes a short adaptation time after transfer to SSM and a high growth rate, likely allowing the mutant to outcompete the original strain during persistent IMI. Thus the original adaptive plasticity comes at a cost.

### The SigB-deficient strain has an increased capacity for proteolysis, casein utilisation, and lactic acid-based fermentation

Casein and lactose are the main protein and main carbohydrate in milk. Strains of varying SigB levels may differ in their ability to metabolise these nutrients. We therefore compare the growth of TG13a, TG13o, and TG13aR in synthetic media containing either casein, lactose, or both (C-medium, L-medium, C+L-medium, respectively). After pre-cultivation in TSB and transfer to the C+L-medium, both SigB-def strains adapt more quickly and have a higher maximum growth rate than the SigB-wt TG13a isolate. Note that these growth patterns are qualitatively similar to those in SSM (Fig. 2A). In the media containing only casein (C-medium) and only lactose (L-medium), the maximum density reached by all strains is much lower, with a slight growth advantage for the SigB-def strains (TG13o and TG13aR) in the C-medium and a comparable disadvantage for these strains in the L-medium. Thus, the SigB-def mutant metabolises casein more efficiently than the SigB-wt in the presence of lactose.

**Fig. 2.**
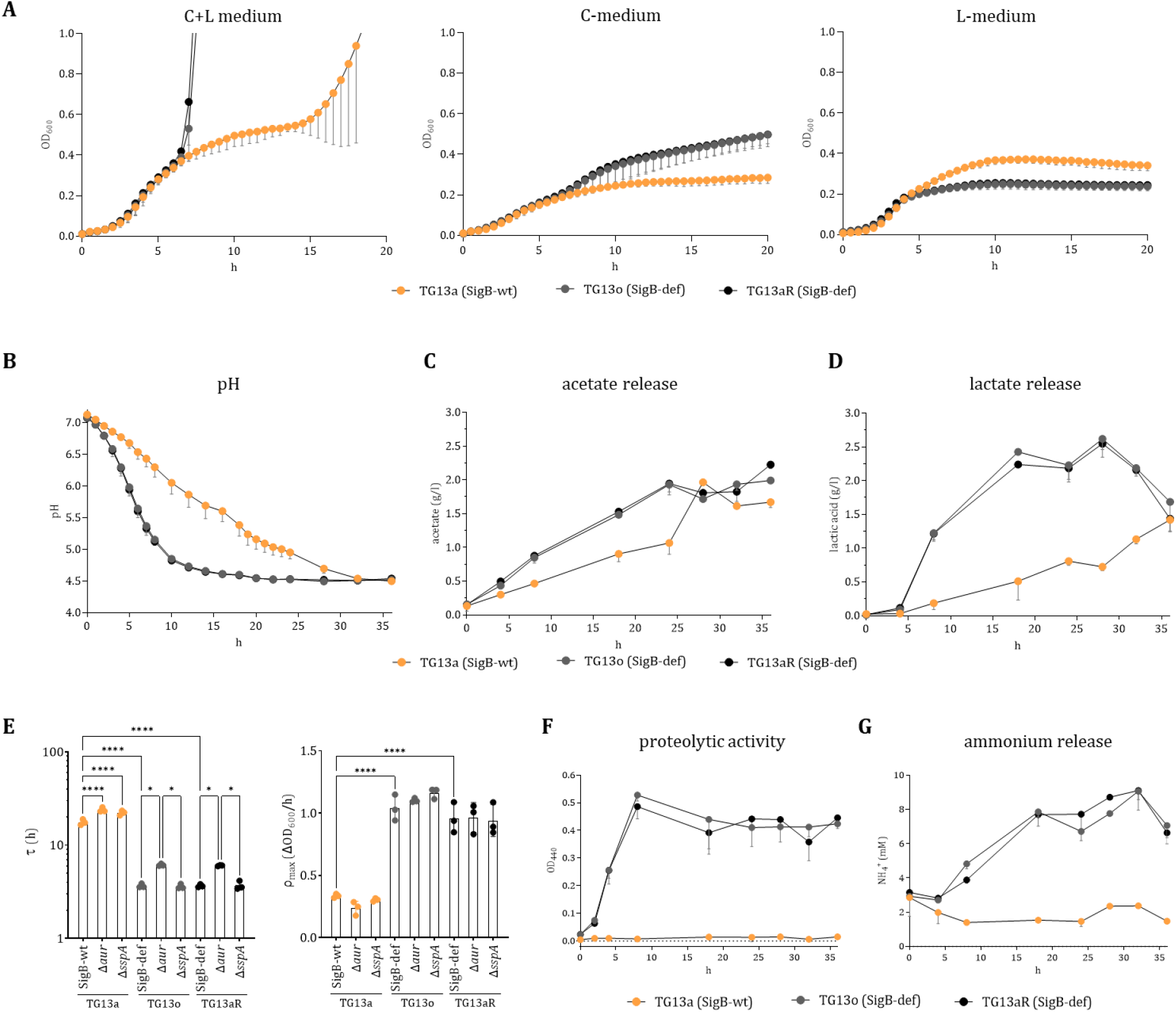
Differences in metabolism between SigB-wt and SigB-def strains. Comparison of TG13a (SigB-wt) and TG13o and TG13aR (both SigB-def). **(A)** Growth curves at OD_600_ (mean ‒ SD) until the end of the experiment or OD_600_ = 1 was reached in media containing (left) casein and lactose (C+L medium), (centre) only casein (C-medium), and (right) only lactose (L-medium). **(B – D)** Dynamics of acidification and associated metabolites from immediately after transfer to SSM up to 36 h later (mean ‒ SD): **(B)** pH, **(C)** acetate release, **(D)** lactate release. **(E – G)** Contribution of proteases and amino acid metabolism: **(E)** Connection between secretion of Aur and SspA and the growth patterns of Δ*aur* /Δ*sspA* isogenic mutants of the SigB-wt (TG13a) and the SigB-def strains (TG13o/TG13aR): maximum growth rate (*ρ*_max_) and adaptation time (*τ*). Linear model analyses were performed on log-transformed data (*τ*) and untransformed data (*ρ*_max_). Differences in pairwise means were tested with Tukey’s HSD. Development of **(F)** proteolytic activity, and **(G)** ammonium Ion (NH_4_^+^) concentration (both mean – SD) across the 36 h window.

Bovine SigB-wt *S. aureus* strains, such as TG13a, have previously been shown to efficiently utilise lactose and cause acidification indicative of fermentation (*3*). We observe a significantly faster decrease in the pH of the SSM media containing the SigB-def strains than in the SSM media containing the SigB-wt strain (Fig. 2B, Supplementary Table S3), as well as a significantly faster accumulation of acetate and lactate (Fig. 2C and 2D, Supplementary Table S3). Interestingly, the SigB-def strains produce mainly lactate as a by-product of fermentation rather than acetate, while the SigB-wt strains produce both of these organic acids at relatively low and comparable levels.

SigB-def strains are known to copiously secrete proteases, particularly the zinc metalloprotease aureolysin (Aur) and serine protease glutamyl endopeptidase (V8 protease; SspA) (*22, 28–30*). These are secreted extracellularly in inactive forms; Aur is capable of not only activating itself but also SspA (*31*). To test the connection between the growth patterns of the SigB-wt (TG13a) and the SigB-def (TG13o, TG13aR) strains in milk and their secretion of Aur and SspA, we conduct experiments using isogenic Δ*aur* and Δ*sspA* knockouts of all strains. The maximum growth rate for both isogenic mutants remained unchanged relative to the original strains. However, Δ*aur* and Δ*sspA* interact epistatically with the SigB-def mutation: in the SigB-def strain, the Δ*aur* knockout significantly prolongs the adaptation time to milk, while the Δ*sspA* knockout has no effect; in the SigB-wt strain, both knockouts show an increased adaptation time, but not to the degree of the single Δ*aur* knockout in the SigB-def strain (Fig. 2E). Generally, bacterial protein degradation can also be monitored directly: in milk, casein aggregates primarily in micelles. Using transmission electron microscopy, we detect collapsed casein micelles in the SSM culture media of the SigB-def strain TG13o (Supplementary Fig. S1C). We also detect substantial casein degradation in the SSM culture media of the SigB-def isolates TG13o and TG13aR and hardly any protein degradation in the SigB-wt (TG13a) culture media (Supplementary Fig. S1D). Correspondingly, proteolytic activity in the culture media of the SigB-def strains increases rapidly within the first 8 h of growth but remains constant at very low levels in that of the SigB-wt strain (Fig. 2F, Supplementary Table S3); ammonia can only be found in the former (Fig. 2G, Supplementary Table S3).

Overall, the SigB-def mutation induces complex metabolic remodelling that is advantageous for growth in milk: SigB-def facilitates the degradation and utilisation of casein and lactose, the main milk components metabolised by *S. aureus*. It further induces metabolic remodelling: lactate rather than acetate becomes the main fermentation by-product of the SigB-def *S. aureus* strains, as in lactic acid bacteria (LAB) (*32*). SigB-def strains also show increased excretion of proteases. Knocking out the aureolysin gene has a stronger effect on the adaptation time in SigB-def strains than in the SigB-wt strain.

### In-host evolution towards SigB-deficiency in bovine mastitis isolates of various clonal complexes

Since SigB-def mutations seem to increase fitness in persistent IMI, we screened 105 consecutively sampled isolates of *S. aureus* from 20 bovine hosts for mutations that disrupt the function of SigB (Supplementary Fig. S2A and B). We find independent SigB-def mutations in three isolates, of which one is assigned to CC151 and two to CC1; the specific mutations differ between isolates (Fig. 3A). As controls, we construct isogenic mutants that reverse two of these mutations. In both reconstructed strains, *asp23* expression is restored, confirming that the mutations are solely responsible for the inactivation of the *sigB* operon (Fig. 3B). All three SigB-def strains show qualitatively and quantitatively similar growth patterns to TG13o: a significantly shorter adaptation time after transfer to milk (Fig. 3C) and a significantly higher growth rate after the initial adaptation phase (Fig. 3D).

**Fig. 3.**
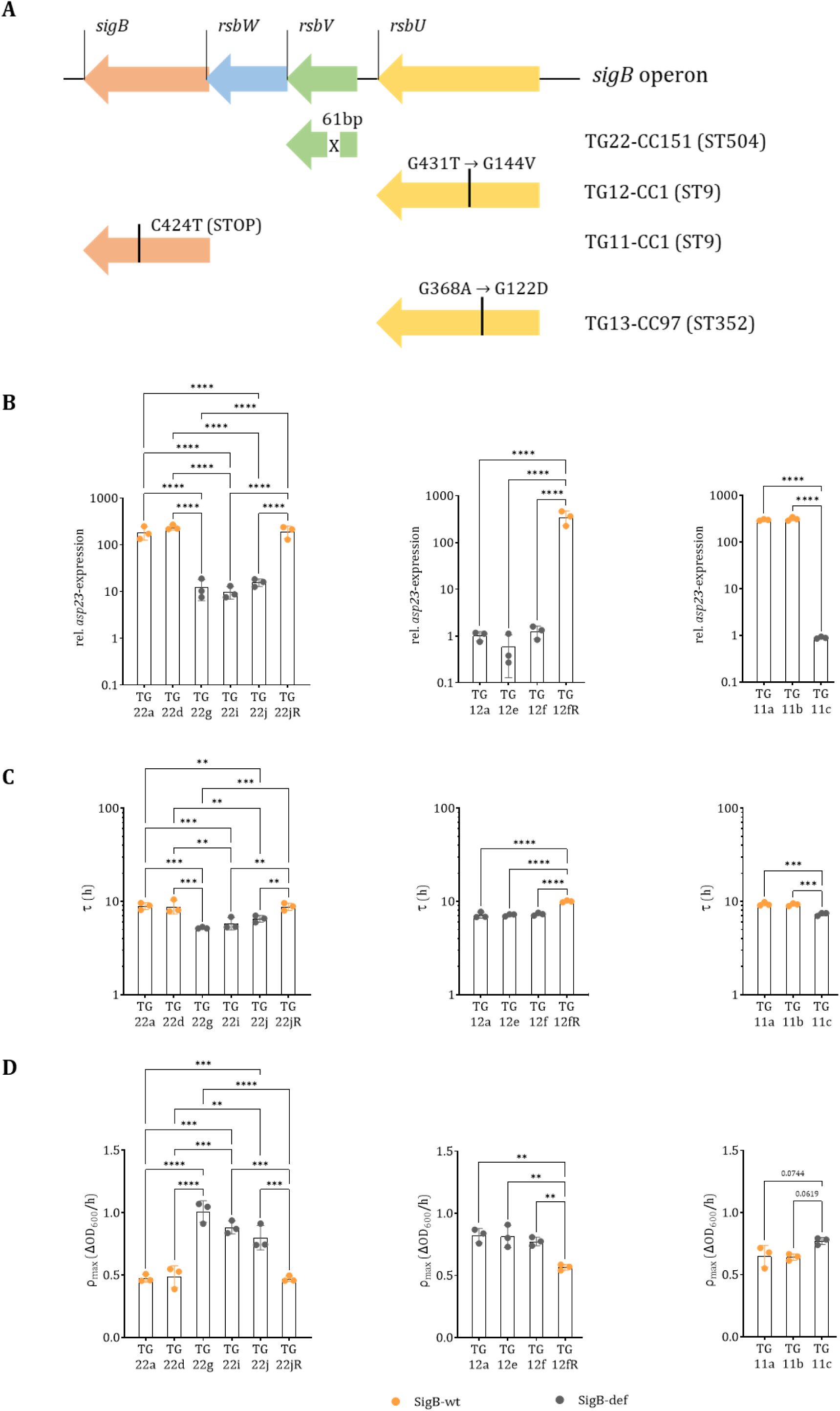
In-host evolution towards SigB-deficiency in bovine mastitis isolates of various clonal complexes. **(A)** Scheme of the *sigB* operon and mutations causing SigB-deficiency that have been identified in *S. aureus* isolates from persistent IMI in the respective isolates. **(B)** Relative *asp23* mRNA expression (RT-qPCR; mean ± SD) after transfer to fresh TSB and harvest at OD_600_ = 3. Differences in pairwise means were tested with Tukey’s HSD. **(C)** Adaptation time (*τ*; mean ± SD) after transfer to SSM. Differences in pairwise means were tested with Tukey’s HSD. **(D)** Adaptation time (*τ*; mean ± SD) and maximum growth rate (*ρ*_max_; mean ± SD) after transfer to SSM. SigB-wt: yellow symbols; SigB-def (different mutations): grey symbols.

Overall, this indicates convergent adaptive evolution of *S. aureus* across clonal complexes. Independent mutations in different parts of the *sigB* operon that reduce SigB activity cause a shorter adaptation time after transfer to milk and a subsequent higher growth rate.

### Niche-specific prevalence of the bovine-associated lineages CC97 and CC151 correlates with SigB activity and growth

Recall that SigB-wt strains of CC97 appear well-adapted to the intracellular niche since they have high adhesion and internalisation capacities and cause only mild IMI with limited tissue damage (*17–19*), while strains of CC151 often cause severe episodes of IMI and more extensive damage (*20, 21*). As these differences may be caused by variation in SigB, we examined a global sample of these two CCs (Supplementary Fig. S3, Supplementary Table S4). We use linear models with clonal complex (CC97 vs CC151) as a fixed effect and isolate as a random effect to show that (i) the CC151 isolates are significantly less pigmented than the CC97 isolates (*p* = 0.0007) and therefore have lower levels of SigB activity, (ii) and that, correspondingly, they have a faster adaptation time and a higher maximum growth rate (both *p*-values < 0.0001) after transfer to SSM. A correlation analysis of all strains, irrespective of their assigned CC, shows a positive relationship between SigB activity (as measured by the level of carotenoid pigmentation) and the adaptation time to SSM, as well as a negative relationship between SigB activity and the maximum growth rate (Fig. 4A).

**Fig. 4.**
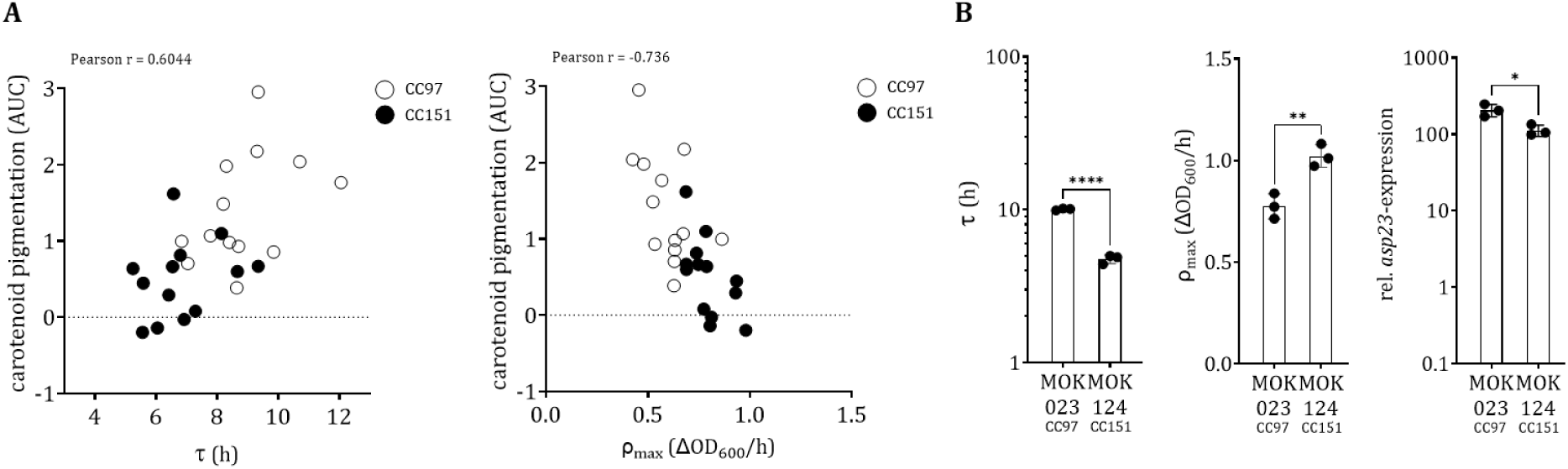
Variation of SigB activity and growth in milk between bovine-associated lineages. **(A)** Carotenoid pigmentation (AUC) positively correlates with the adaptation time (*τ*), and negatively correlates with the maximum growth rate (*ρ*_max_) of CC151 (n = 13) and CC97 (n = 13) isolates. Each point represents one isolate of CC151 (black) and CC97 (white). **(B)** Comparison of MOK023-CC97 and MOK124-CC151 after transfer to SSM. Differences in mean were evaluated using t-tests of log-transformed data for relative *asp23* mRNA expression and for adaptation time (*τ*) and of untransformed data for maximum growth rate (*ρ*_max_), respectively.

We corroborate these results by separately analysing a CC97/CC151 strain pair that has been clinically well-characterised *in vitro* and via infection experiments (*18, 19, 21*): Indeed, the *in vitro* growth phenotypes for MOK023-CC97 and MOK124-CC151 strains mimic those observed *in vivo* immediately after onset of the bovine intramammary challenge experiment (*21*). MOK124-CC151 has a significantly shorter adaptation time than MOK23-CC97 and a significantly higher maximum growth rate. Note that both strains are SigB-wt but differ significantly in their *asp23* transcription levels, with MOK124-CC151 specifically showing reduced SigB activity (Fig. 4B).

Overall, differences in SigB activity between SigB-wt strains of different CCs allow them to more effectively adapt to and subsequently survive in specific niches of the bovine udder.

### Human-associated *S. aureus* strains with SigB-def mutations adapt better to milk than otherwise identical SigB-wt strains

If low SigB activity generally enhances growth in milk, then human-derived strains - the major source of newly emerged bovine-adapted strains (*33, 34*) - could similarly exhibit enhanced growth if they are SigB deficient. Indeed, human-associated SigB-def strains of the globally, well-distributed CC8 lineage (*35*) (both isolated and engineered) grow vigorously after about 15 h while the otherwise genetically identical SigB-wt strains grow only much later and more slowly if at all (Fig. 5A, Supplementary Table S5). Note that the adaptation times for the SigB-def human-isolated strains are actually long compared to even the SigB-wt bovine strains. This is expected since human-isolated strains may not have encountered a milk-rich niche during their evolution.

**Fig. 5.**
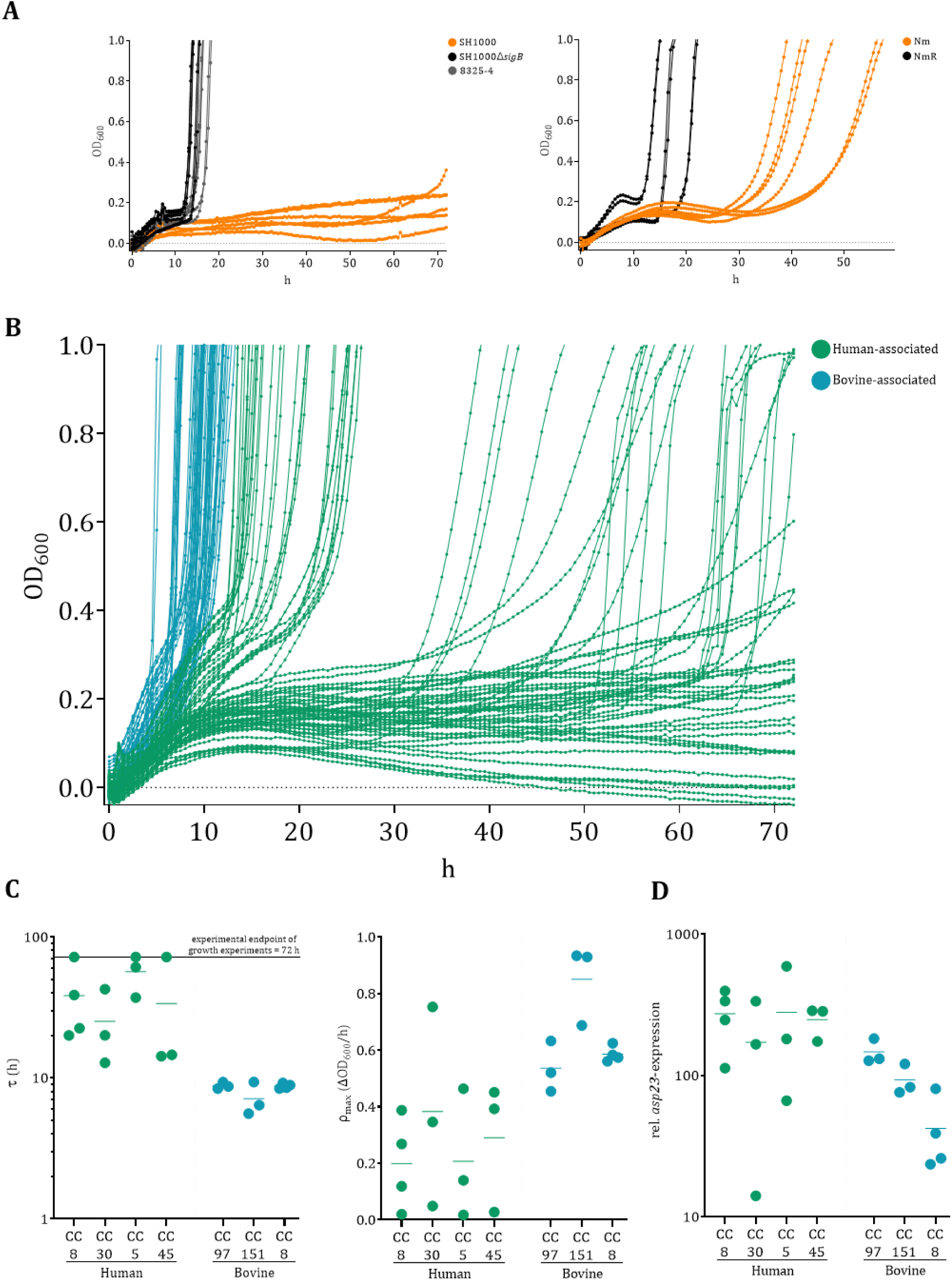
Variation of SigB activity and growth in milk between and among human- and bovine-associated isolates. **(A and B)** Growth dynamics after transfer to SSM are shown until an OD_600_ = 1.0 is reached or until the end of the observation period. For each strain, growth curves of three independent experiments comprising two technical replicates are shown: **(A)** After transfer to SSM, human-associated SigB-wt strains grow late and slowly while the isogenic SigB-def strains grow faster. Shown here are the human-derived SigB-def strain 8325-4 (grey), the repaired SigB-wt mutant SH1000 (orange), and the engineered SigB-def mutant SH1000Δ*sigB* (black) as well as the human-derived Newman strain Nm (SigB-wt, orange) and the isogenic SigB-def mutant NmR (black). **(B)** Bovine-associated strains (blue) grow faster than human-associated strains (green), irrespective of the CC. **(C)** Adaptation time (*τ*) and maximum growth rate (*ρ*_max_) after transfer to SSM. The adaptation time was set to 72 h for three human isolates because they did not reach an OD = 0.45 after 72 h or did not grow at all. **(D)** SigB activity determined by relative *asp23* mRNA expression (RT-qPCR) after strains were grown to an OD_600_ = 3 in TSB.

Overall, human-derived strains that grow slowly or not at all in milk can adapt to growth in milk through deletion of SigB.

### Bovine-associated *S. aureus* strains grow much better in milk than human-associated strains

We now extend our analysis of the growth patterns of human SigB-wt isolates by including strains from CC5, CC30, and CC45 in addition to strains from CC8 (*35*), and can confirm that they all grow slowly in milk if at all (Fig. 5B, Supplementary Table S6). By comparison, the bovine *S. aureus* strains assigned to CC8, CC97, and CC151 all grow well when transferred to SSM (see Fig. 5C for adaptation time; *p* = 0.0320 and for maximum growth rate; *p* = 0.0382). Using linear models, where we excluded the bovine-associated CC8 strain because it only recently host-jumped to cows (*36, 37*), we find that while expression of *asp23* is lower on average in the bovine strains, the difference is not significant (*p* = 0.2308) (Fig. 5D). This indicates that SigB activity is not the sole driver of the increased growth rates of bovine strains and additional metabolic adaptations may be involved. However, when CC8 is considered separately, we do observe lower SigB activity in the bovine strains (*p* = 0.0004) that matches their faster adaptation time (*p* < 0.0001) and higher maximum growth rate (*p* = 0.0005).

Overall, SigB activity does not differ between human and bovine strains or CCs, although the bovine strains adapt better to milk. However, in the case of CC8, where human and bovine isolates are particularly closely related, we do observe lower SigB activity levels in the bovine strains that matches their faster growth in milk. Therefore, modulation of SigB activity may initiate a broader metabolic remodelling process.

### A transposon mutant library screen shows that the *sigB* operon is the staphylococcal transcriptional regulator with the shortest adaptation time to milk

Such a metabolic remodelling process could be driven by a diverse set of transcription factors. We therefore analyse the growth patterns and proteolytic activity of all the mutant strains in the Nebraska transposon mutant library (NTML) (*38*), which comprises strains with disrupting mutations in transcriptional regulator genes, i.e., sigma factors and transcription factors (n = 103; Supplementary Table S7) (*39*). The original JE2 *S. aureus* strain that forms the reference of the library was derived from a human CC8 isolate; as expected, JE2 adapts and grows relatively slowly after transfer to SSM. The *sigB* mutant has the shortest adaptation time of all mutant strains and a comparatively high maximum growth rate (Fig. 6A and B, and Fig. 7A).

**Fig. 6.**
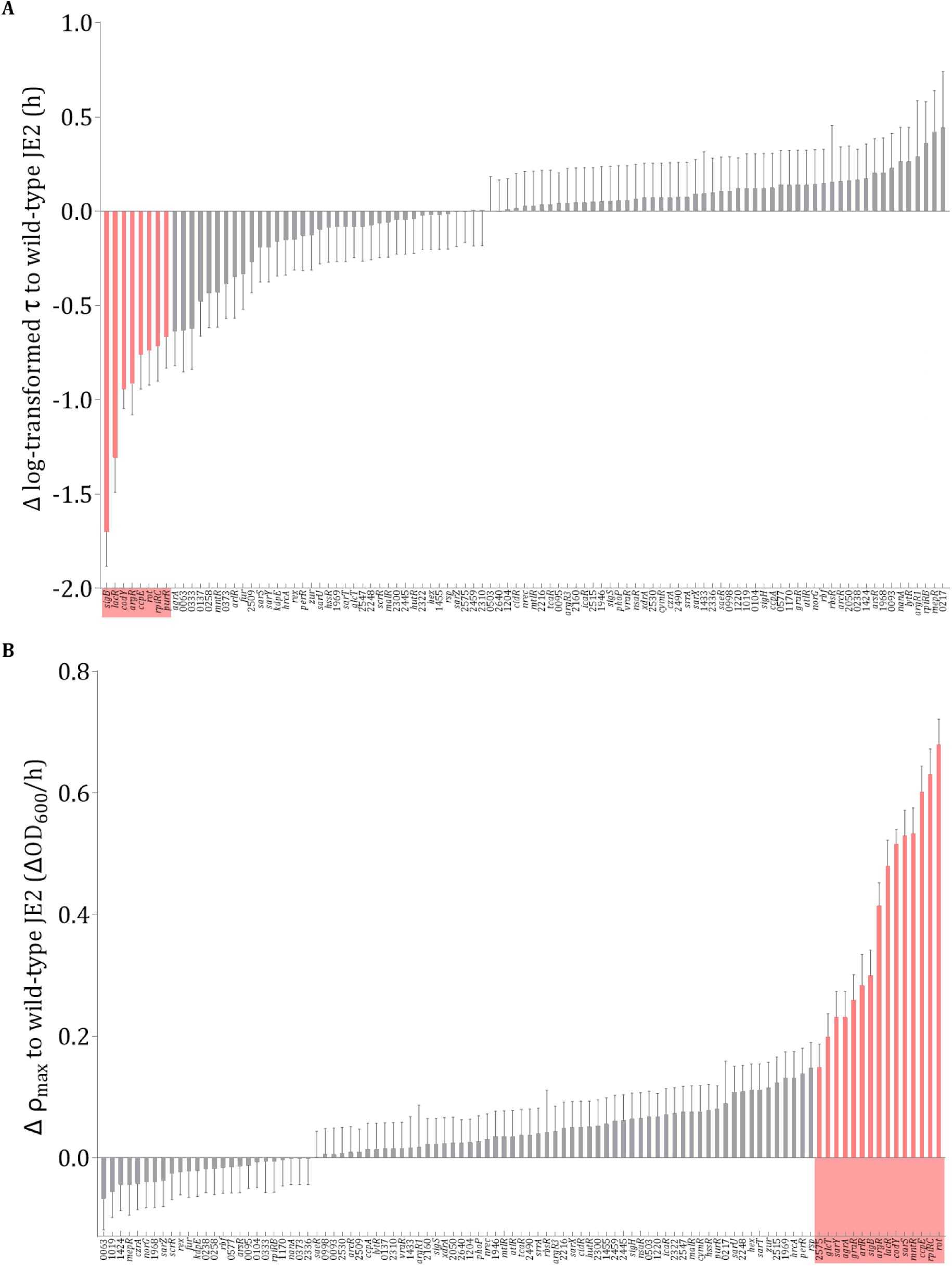
Screening of NTML regulatory mutant strains (part I). Means and standard errors of the differences between each NTML mutant strain and the wild-type reference strain with respect to **(A)** adaptation time (*τ*) and **(B)** maximal growth rate (*ρ*_max_). The pink colour indicates significant differences as determined by Dunnett’s test with the wild-type as a reference. The four mutants *gltC*, *treR*, *sarA,* and NE40 (SAUSA300_1174) were excluded from the analysis because they did not grow in SSM within 72 h.

**Fig. 7.**
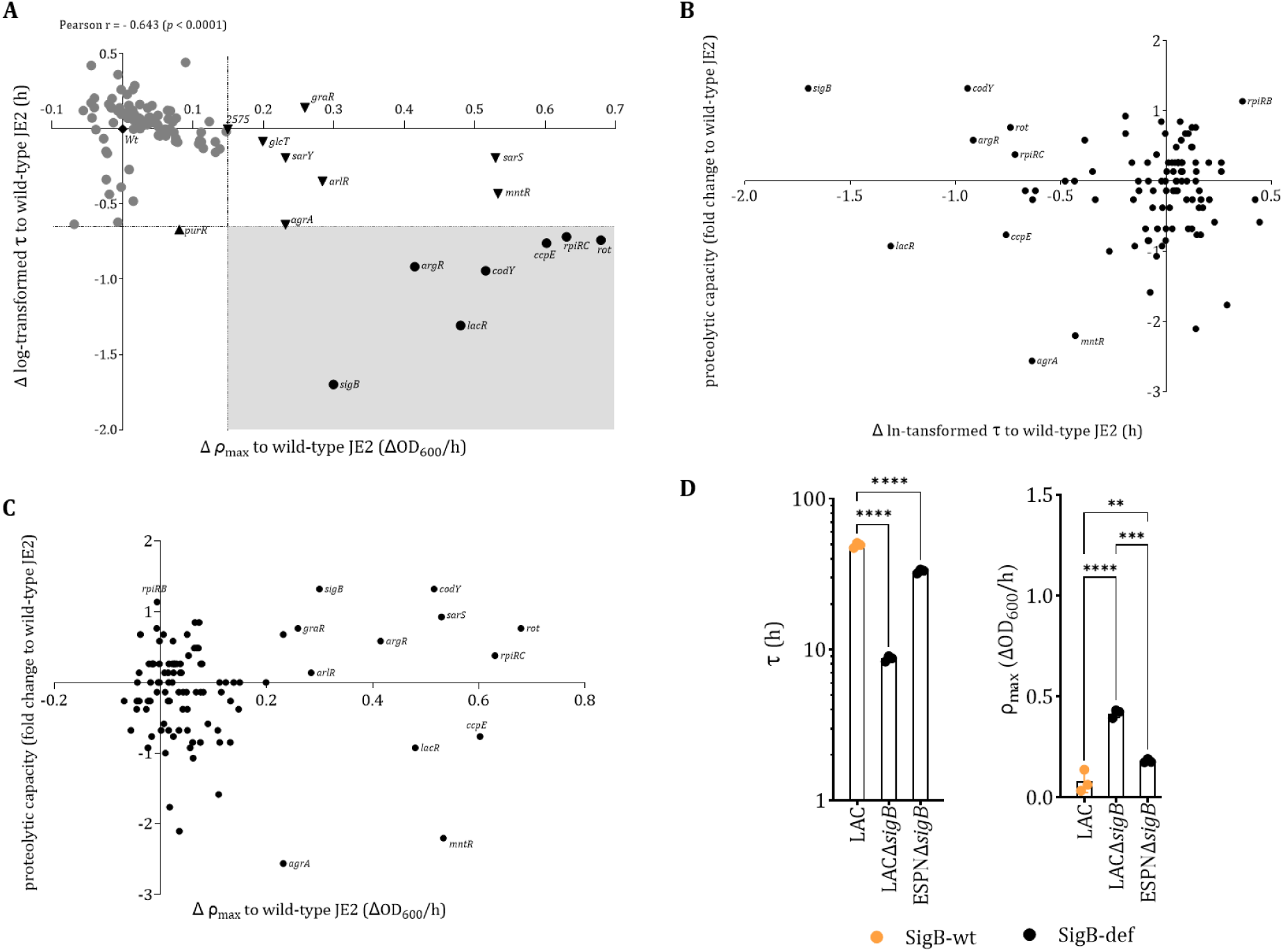
Screening of NTML regulatory mutant strains (part II). **(A)** Scatter plot and correlation of the differences in the log-transformed adaptation time (*τ*) and in the maximum growth rate (*ρ*_max_) between the 103 NTML regulatory mutant strains and the original JE2 strain. Seven mutants exhibiting a significantly shorter adaptation time as well as a significantly higher maximum growth rate than the JE2 strain are underlaid with a grey rectangle. The four mutants *gltC*, *treR*, *sarA,* and NE40 (SAUSA300_1174) were excluded from the analysis because they did not grow in SSM within 72 h. For each NTML regulatory mutant strain, the difference in proteolytic capacity from the reference JE2 strain (on the x-axis) is plotted **(B)** vs the difference in the ln-transformed adaptation time (*τ*), and **(C)** the maximal growth rate (*ρ*_max_) (y-axis) from the reference JE2 strain. The pairwise Pearson correlations are not significant and therefore not shown. **(D)** Impact of protease-null mutants on *S. aureus* growth in a SigB-def background on the adaptation time (*τ*), and the maximum growth rate (*ρ*_max_). An ANOVA on log-transformed data was performed for adaptation time (*τ*) and on untransformed data for the maximum growth rate (*ρ*_max_); in both cases, differences in mean were tested with Tukey’s HSD.

Although casein is the most important source for amino acids in milk and can be efficiently utilised by SigB-def strains, the proteolytic activity of *S. aureus* strains surprisingly does not appear to influence their adaptation time or growth rate in this screen (Fig. 7B and 7C). While loss of SigB activity induces faster growth in SSM, the growth phenotype is only partially reverted to the wild-type level when all staphylococcal proteases are absent (Δ*sigB*Δ*aur*Δ*sspAB*Δ*scpA*Δ*spl*) (Fig. 7D). Thus the loss of SigB causes further not yet fully investigated metabolic remodelling that is favourable for growth in milk, independent of increasing the proteolytic activity. Similar remodelling is induced by knockouts of the negative regulator of the lactose utilisation reagulator *lacR*, of the regulator of the branched-chain amino acid synthesis *codY*, and of the regulator of arginine biosynthesis *argR*, which are among the seven mutants with significantly decreased adaptation times and increased growth rates (see Fig. 7A).

Overall, low SigB-activity seems to “pre-adapt” even human-associated strains so that they grow well in milk. This is evidenced by the *sigB* mutant having the shortest adaptation time of all mutants that disrupt the functions of the transcriptional regulator genes of *S. aureus*, as well as a comparatively high maximum growth rate. *S. aureus* proteolytic activity only partially contribute to the elevated growth phenotype in milk, alongside significant metabolic remodelling in SigB-deficient strains.

## Discussion

Throughout this article, we employ an *in vitro* system where strains of *S. aureus* that have been maintained in the standard laboratory medium tryptone soy broth (TSB) are switched to semi-skimmed milk broth (SSM) and their adaptation time and maximum growth rate in SSM are subsequently monitored. With this approach, we show that the phenotypically plastic SigB-wt strain TG13a, isolated early during a persistent IMI and assigned to CC97, can adapt well to milk. However, the SigB-def mutant strain TG13o, which evolved later during IMI and completely replaced its ancestor, has an even shorter adaptation time and a higher growth rate in milk. Genetic engineering confirms that the SigB-def mutation causes this growth advantage and is associated with broader metabolic remodelling: SigB-def leads to reduced carotenoid pigmentation and expression of adhesins (traits that promote adaptation to the intracellular niche) and to degradation and utilisation of the main components of milk, casein and lactose (traits that promote adaptation to the extracellular niche). In a competition experiment, the SigB-def strain outgrows the SigB-wt strain after a switch from TSB to SSM, while neither strain has a consistent advantage in TSB. Other *S. aureus* strains isolated from persistent IMI have evolved repeatedly from SigB-wt towards SigB-def with similar phenotypic consequences, irrespective of their CC. The evolution of SigB-def strains with canalised phenotypes that are narrowly adapted to milk from SigB-wt strains with plastic phenotypes that are capable of expressing carotenoid pigments and adhesins and therefore exploiting an intracellular niche (*5, 6, 11*), constitutes a case of genetic assimilation. This genetic assimilation appears driven by the cost of phenotypic plasticity: longer time-lags in adapting to milk and subsequent slow growth. Analogously, persistent intracellular infections may lead to fixation of canalised SigB-co mutations, since long-term intracellular passage of *S. aureus* appears to favour high SigB activity and this may be genetically fixed (*11, 25*) (Fig. 8).

**Fig. 8.**
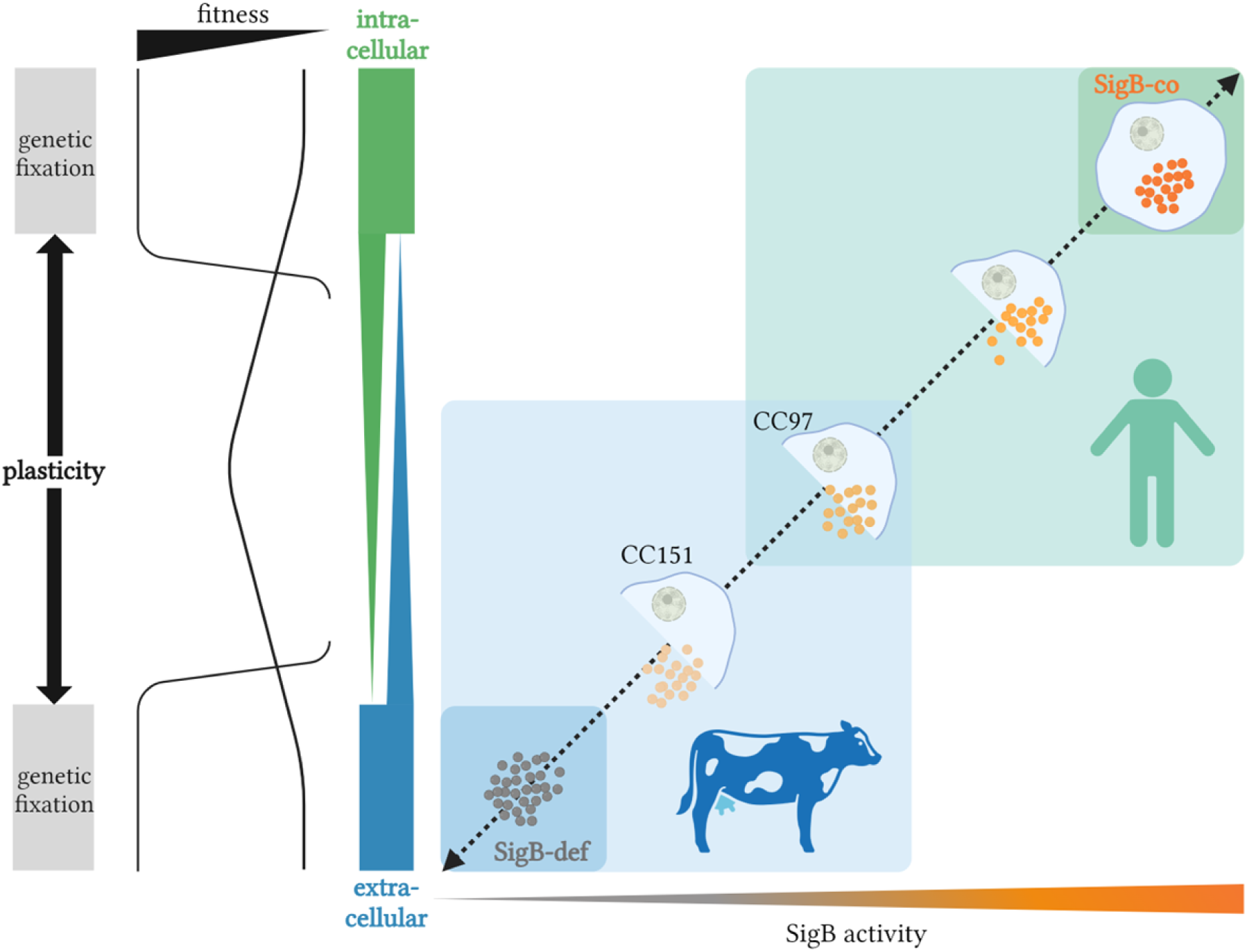
Plastic and genetic regulation of SigB activity governs a large part of *S. aureus* niche adaptation. Model of the divergent roles of SigB in the adaptation of *S. aureus* to intra- and extracellular niches showing the connections between the SigB genotype and carotenoid pigmentation (a proxy of SigB activity and a known adaptation to the intracellular niche). High SigB activity induces traits necessary for intracellular survival and growth; low SigB activity induces traits for extracellular growth. The ability to switch between niches seems to require a plastic SigB-wt state. At both ends of the spectrum, SigB-wt strains are outcompeted by SigB-def or SigB-co strains. SigB-wt bovine isolates of CC97 generally exhibit relatively high levels of SigB and have lower fitness in milk than SigB-wt bovine isolates of CC151. Overall, bovine-associated isolates adapt more readily to milk than human-associated isolates. Created in https://BioRender.com.

The two main bovine clonal complexes of *S. aureus* are CC97 and CC151. SigB-wt strains of CC97 cause subclinical IMI and seem to be particularly well-adapted to the intracellular niche, while SigB-wt strains of CC151 cause episodes of clinical IMI and seem better adapted to the extracellular niche (*17–21*). These differences in clinical severity and niche-specificity are consistent with the results obtained from comparison of SigB-def mutant strains with their SigB-wt isogenic progenitors in our *in vitro* system: lower levels of SigB activity in CC151 compared to CC97 correlate with shorter adaptation times and faster growth. Thus SigB activity seems to both quantitatively and qualitatively regulate the niche use of bovine-adapted strains: subtle shifts in the phenotypically plastic strains are quantitative changes, and loss-of-function mutations to a canalised phenotype are qualitative changes (Fig. 8). Notwithstanding the adaptive potential of mobile genetic elements (*3, 40*), adaptive plasticity and evolution mediated by SigB, a core gene and master regulator of transcription, is crucial for niche adaptation in *S. aureus*.

SigB-wt strains isolated from humans have likely never encountered a milk-rich extracellular niche such as the bovine udder during their evolution. As expected, they generally grow slowly when switched to milk, if at all. Nevertheless, the isogenic SigB-def counterparts of these human-associated SigB-wt strains adapt relatively quickly to milk and then grow well. Thus, the SigB-def mutation seems to “pre-adapt” (*41*) human strains to a milk-rich environment. Indeed, SigB-def mutants in *S. aureus* may facilitate adaptation to extracellular niches in general rather to only milk-rich ones: clinical isolates from persistent human cystic fibrosis (CF) infections are often characterised by SigB-def mutations(*42–45*).

As in *S. aureus,* modulation of SigB activity is crucial for the transition between intra- and extracellular niches in *L. monocytogenes* (*46*). However, in contrast to the adaptation process in *S. aureus*, expression levels of SigB are high in the extracellular host niche and low in the intracellular niche (*47*). This reversal of the regulatory mechanism of SigB indicates that its specific role in niche adaptation in both species evolved independently from the ancestral role of stress response coordination.

Importantly, remodelling of metabolic processes that are not regulated by SigB also appear to contribute to host-specific adaptation: Although there is no significant difference in the average SigB activity of bovine and human SigB-wt isolates (as measured via proxies of SigB activity), all bovine isolates adapt better to milk than human isolates. Screening a transposon mutant library, we identify the regulators of some of these metabolic processes: the negative regulator of lactose utilisation LacR, as well as the regulators of the branched-chain amino acid synthesis CodY and arginine biosynthesis ArgR. Their pathways have previously been identified as important for growth in milk using a multi-omics approach (*48*).

The evolution of the canalised SigB-def mutations from the plastic SigB-wt constitutes a case of genetic assimilation. Analogously, persistent intracellular infections may lead to fixation of canalised SigB-co mutations, since long-term intracellular passage of *S. aureus* appears to favour high SigB activity that may be genetically fixed (*11, 25*). Thus, canalisation of high or low SigB activity levels when *S. aureus* persists long enough in exclusively intra- or extracellular niches may be a general phenomenon. However, phenotypic plasticity and therefore the ability to switch between intra- and extracellular niches may be necessary to infect new hosts. Instead of qualitative changes via SigB-def or SigB-co mutations, maintenance of adaptive plasticity and quantitative, polygenic modulation of SigB activity (*46*) may ensure long-term survival and continued adaptability.

## Materials and Methods

### Strains

All strains were stored in glycerol stocks at -80 °C and were pre-cultivated on tryptic soy agar (TSA) (Thermo Fisher Scientific, Oxoid, Hampshire, UK) at 37 °C for 24 h and kept at 4 °C for a maximum of two weeks until use. TSA for strains of the Nebraska Transposon Mutant Library (NTML) and the SH1000Δ*sigB* mutant was supplemented with erythromycin (5 µg/mL) (Carl Roth, Karlsruhe, Germany), and TSA for LACΔ*sigB* and LAC* protease KOΔ*sigB* mutants was supplemented with tetracycline (5 µg/ml) (Carl Roth, Karlsruhe, Germany).

#### Bovine-associated isolates from persistent IMI and genetically engineered mutants

The isolates TG13a (SigB-wt) and TG13o (SigB-def) were described in our previous study (*22*). The single nucleotide change *rsbU*(G368A) observed in TG13o was introduced into TG13a, and the resulting strain designated as TG13aR (*24*). In addition, we engineered isogenic knockout mutants of the proteases Aur (Δ*aur*) and SspA (Δ*sspA*) based on the SigB-wt (TG13a) and SigB-def strains (TG13o/ TG13aR).

The isolates TG11-CC1 (n = 3), TG12-CC1 (n = 3) and TG22-CC151 (n = 5) were obtained by retrospectively screening 105 staphylococcal isolates from the milk of 20 cows with persistent IMI for visible loss of colony pigmentation (i.e., for a white or grey instead of a yellow colour). Note that we consider IMI persistent when clonal isolates can be collected from at least three consecutive milk samples from the same dairy cow, with a minimum of one month between the first and the last of these three samples. This screen yielded putative SigB-def isolates in milk samples from three cows from two herds. The isolates TG22-CC151 were from one udder quarter of a single cow (cow 11 in study Grunert, et al. 2018): Isolates from samples taken within the first two weeks (TG22a and d) are SigB-wt, while those from samples taken in weeks 33 (TG22g), 52 (TG22i), and 64 (TG22j) are SigB-def. The isolates TG11-CC1 (n = 3) and TG12-CC1 (n = 3) came from cows of the same dairy herd in Austria, and were collected from milk samples taken during routine microbial mastitis diagnostic screening over 15 weeks. In fact, TG11-CC1 and TG12-CC1 isolates were collected at weeks 1, 4 and 15 from two different cows. While TG11a and TG11b are SigB-wt, a stop codon in the *sigB* gene of TG11c causes SigB-deficiency. All three TG12 isolates (a, e, and f) are SigB-def due to the SNP G431T in *rsbU*. Genome sequencing of these isolates reveal clonality within isolates from the same cow, indicating persistent infection with the same strain. Although the isolates TG11 and TG12 were recovered from different cows, genome sequencing indicates clonality, suggesting transmission from one cow to the other. We also engineered the following isogenic mutant strains: In strain TG22jR, we repaired the 62 bp truncation in *rsbV* in the last collected isolate (TG22j), and in strain TG12fR we reversed the point mutation *rsbU*:431G→T (*24*).

The characteristics of all isolates are summarised in the Supplementary Table S1 and Supplementary Fig. S2C and D. The procedures followed to engineer the mutants are described in the subsection Genetic manipulation.

#### Bovine isolates for the comparison of CC97 and CC151

We selected the strain pair MOK023-CC97 and MOK124-CC151 because comprehensive *in vivo* and *in vitro* data are available (*18, 19, 21*). MOK023-CC97 (ST3170) and MOK124-CC151 (ST151) strains were originally isolated from cases of clinical mastitis (*49*). The CC97 and CC151 isolates (n = 13, each) used for comparative analyses in this article originate from our collection of well-characterised strains from bovine mastitis milk samples (*50–57*). Strains were selected to represent diverse geographic origins (Austria, n = 11; Argentina, n = 9; Rwanda, n = 4; others, n = 2), sequence types (STs), and *spa*-types. Core genome multilocus sequence typing (cgMLST) and subsequent analyses are described in the subsection: Genome sequencing and molecular genetic typing. A comprehensive overview of the geographic origin, STs, *spa*-types and cgMLST analyses of all strains is provided in Supplementary Table S4 and Supplementary Fig. S3.

#### Human-associated isolates and genetically engineered mutants

Strain 8325-4 is a widely used model strain originally isolated from a sepsis patient in 1960 (strain RN1 or NCTC8325); all three prophages were cured by UV radiation (*58*). Strain 8325-4 is SigB-def because of an 11-bp deletion in *rsbU*, which was repaired in strain SH1000 (SigB-wt) (*28*). The isogenic mutant SH1000Δ*sigB* (SigB-def) was derived from SH1000 (*11*). Strain Newman (Nm, NCTC8178) is SigB-wt and was isolated from a patient with osteomyelitis (*59*). We introduced the same single nucleotide change in *rsbU*(G368A) that we studied in TG13o to the Newman strain to make the SigB-def mutant strain NmR (*24*). The SigB-wt strain LAC (CA-MRSA USA300) was originally isolated from a human skin and soft tissue infection (*60*), and we engineered the isogenic mutant LACΔ*sigB* (SigB-def) from it. In addition, we combined a derivative of the LAC strain that lacks all 10 major secreted proteases (LAC*protease KO) (*61*) with a SigB-def mutant, LAC* protease KOΔ*sigB*. The characteristics of all isolates are summarised in Supplementary Table S5, and the procedures followed to engineer the mutants are described in the subsection: Genetic manipulation.

#### S. aureus isolates for comparison of human- and bovine-associated CCs

A set of 13 human- and 10 bovine-associated isolates was used. The subset of human isolates includes major MRSA and MSSA epidemic clones, specifically: Four isolates of CC8, and three isolates of CC30, CC5, and CC45 each. These represent CCs commonly isolated from asymptomatic and infected humans (*59, 62, 63*). The subset of bovine-associated isolates comprises three strains each of CC97 and CC151, and four isolates of bovine CC8 (*51, 54–57*). Within each CC, strains were selected to represent diverse sequence types (ST, MLST) and *spa*-types.

All isolate characteristics are summarised in Supplementary Table S6.

#### S. aureus Nebraska Transposon Mutant Library (NTML)

The Nebraska Transposon Mutant Library (NTML) provides a comprehensive collection of knockouts of specific genes from a wild-type strain, including knockouts of transcriptional regulators, via transposon mutagenesis (*38*). The wild-type strain JE2 is a USA300 plasmid-cured derivative (USA300_FPR3757) of strain LAC (*60*), a well-characterised community-associated methicillin-resistant *S. aureus* (CA-MRSA) strain isolated from the Los Angeles County jail. Plasmids were cured to facilitate genetic manipulation and avoid interference during transposon mutagenesis experiments, resulting in strain USA300 JE2. A list of 108 NTML strains, where transcriptional regulators are knocked out, is provided in Gimza *et al.* 2019. Of these 108 mutants, 103 were available at the time of acquisition of the NTML (Supplementary Table S7).

#### Comparison with data from Gimza et al

Gimza *et al.* 2019 screened transposon mutants for all 108 available transcriptional regulators in *S. aureus* USA300 JE2 from the Nebraska Transposon Mutant Library (NTML) and reported the alterations in proteolytic activity using gelatin zymography normalised to the band intensity of the USA300 JE2 wild-type strain (Supplementary Table S7). We extracted the fold change in band intensity from their Fig. 5, transformed it to the log_2_ fold ratio between mutant and wild-type, and performed a pairwise Pearson correlation analysis of the proteolytic capacity and adaptation time τ, and of the proteolytic capacity and maximum growth rate *ρ*_max_ (Fig. 7B and C).

### Genetic manipulation

The thermosensitive plasmid pIMAY-Z (*64*) was utilised to generate *aur* and *sspA* deletion mutants using previously described protocols (*65*). Briefly, oligonucleotide primers were used to amplify ∼500 bp AB and CD flanking regions of each gene of interest (Supplementary Table S8) using Q5 polymerase (NEB). Gibson assembly (NEB) generated pIMAY-Z constructs that were transformed firstly into *E. coli* DC10B (*65*) and then, after Sanger sequencing confirmation (Eurofins), into the *S. aureus* strain of interest. After integration at 37°C and plasmid excision at 30°, OUT PCR was used to confirm that the clones contained gene deletions.

To restore the truncation in *rsbV* TG22j, the full-length allele (including AB and CD flanking regions) was amplified (Supplementary Table S8) from the precursor isolate TG22a and cloned into pIMAY. Allelic exchange was performed as described above. The resulting strain was named TG22jR.

The protease null strain (Δ*aur*Δ*sspAB*Δ*scpAspl*::erm) was originally generated by Wörmann et al. (*61*) and the originating *sigB*::tet mutation was generated by Chan, et al. (*66*). Strains were constructed using Φ11 phage lysates containing *sigB*::tet and transduced into either the USA300 LAC background (*67*) or the USA300 LAC protease null background. Transduced strains were selected on TSA containing 5 µg/mL tetracycline and confirmed via PCR using the primers OL7735 and OL7736.

The oligonucleotides employed for genetic manipulation are summarised in Supplementary Table S8.

### Genome sequencing and molecular genetic typing

Genomic DNA for whole-genome sequencing (WGS) was extracted from the *S. aureus* isolates using the MagAttract HMW DNA Kit (Qiagen, Hilden, Germany) according to the manufacturer’s instructions. DNA concentration was checked using a Qubit 3.0 fluorometer (Thermo Fisher Scientific, Waltham, MA, USA) and the dsDNA HS assay kit (Thermo Fisher Scientific). The library was prepared with the Nextera XT kit (Illumina Inc., San Diego, CA, USA), followed by paired-end sequencing (2 x 300 bp) on an Illumina MiSeq device. The quality of the raw reads was checked using FastQC v0.11.9. Adapter sequences were trimmed with Trimmomatic v0.36 using default parameters (*68*). Afterwards, the genome was assembled using SPAdes v3.11.1, and contigs were filtered by requiring a minimum coverage of 5x and a minimum length of 200 bp (*69*). Conventional multilocus sequence typing (MLST) and core genome (cg)MLST were performed with the Ridom SeqSphere+ software v.9.0.8 (Ridom, Münster, Germany). For the latter, we used the official scheme comprising 1861 target genes available in that software (https://www.cgmlst.org/ncs/schema/schema/141106/). A minimum spanning tree (MST) was then generated to visualise clusters and the genetic relatedness between the isolates, using a cluster threshold of 24 alleles. Antimicrobial Resistance Genes (ARGs) and point mutations conferring antibiotic resistance were extracted with ResFinder. Plasmid replicons were extracted with PlasmidFinder (*70, 71*).

Locus-specific sequencing of the *sigB* operon (*rsbU*-*rsbV*-*rsbW*-*sigB*) was performed as previously described (*24*). MLST was conducted following Enright, et al. (*72*), except that we adjusted the trimming position for defining *gmk* alleles to 417 bp instead of 429 bp. Allelic profiles and ST were determined using the PubMLST website at https://pubmlst.org/organisms/staphylococcus-aureus. For *spa*-typing, the sequence of a polymorphic VNTR located in the 3’ coding region of the *S. aureus*-specific staphylococcal protein A (*spa*) was utilised (www.spaserver.ridom.de) (*73*). *Spa*-types were categorized based on the repeat sequence using the spa typer application in the Ridom SeqSphere+ software v.9.0.8 (Ridom, Münster, Germany). *Agr*-groups were identified through multiple PCR techniques as outlined by Gilot, et al. (*74*).

### Growth media

Unless otherwise indicated, we used semi-skimmed milk (SSM) broth media, which consisted of a 1% w/v mixture of skimmed milk powder (LP0031, Thermo Fisher Scientific, Oxoid, Hampshire, UK) in distilled water, equivalent to 10% fresh milk with ≤1.5 % w/w fat content. SSM was sterilised by autoclaving at 121°C for 5 min. Tryptone Soy Broth (TSB) was used for overnight cultures and growth control: it consisted of 30 g of powder (CM0129B, Thermo Fisher Scientific, Oxoid, Hampshire, UK) dissolved in 1 litre of distilled water and sterilised by autoclaving at 121°C for 15 min. Bacteria were grown in whole milk, specifically in 1 % v/v commercially available fresh dairy milk (3.5 % w/w fat content) diluted in distilled water. To study the influence of casein and lactose, the base minimal medium Roswell Park Memorial Institute Medium (RPMI-1640 without glucose; Sigma Aldrich, Merck, Germany) was supplemented either with 1 % casein sodium salt (Sigma Aldrich) to obtain the C-medium, or with 1 % D-Lactose monohydrate (Sigma Aldrich) to obtain the L-medium, or an equal mixture of 1% casein and lactose to obtain the C+L-medium.

### Growth measurements

Bacteria were grown overnight (17 - 19 h) at 37 °C in 3 ml TSB (preculture). Cell density was measured at 600 nm (OD_600_) in a cuvette using a photometer (Eppendorf, Hamburg, Germany). Bacterial cultures were diluted 1:5 in RPMI-1640 medium (Sigma-Aldrich) to minimise bacterial growth during handling and the transfer of the original TSB medium to the next growth medium. For the growth measurements taken in SSM, diluted bacterial cultures (approx. 10 µl) were used to inoculate SSM in a microplate, specifically 200 μl SSM per well; the final starting point for the experimental culture was OD_600_ = 0.05. In addition, diluted bacterial cultures were always transferred to fresh TSB as a parallel control. Plates were grown at 37 °C without shaking for 72 h. Cell growth was automatically measured at OD_600_ every 30 min using Bioscreen-C (Oy Growth Curves Ab Ltd, Helsinki, Finland) after pre-shaking at medium intensity for 10 s. The growth curve obtained from the medium with bacteria (OD_600_) was subtracted from records of the respective media without bacteria (blank at OD_600_) over the entire time course (72 h). Note that this subtraction may result in negative measurements when bacterial concentrations are low, especially at early time points. We used the same procedure to screen whole milk in 1 % v/v, C-medium, L-medium, and C+L-medium using a plate reader.

In SSM, growth of the *S. aureus* strains TG13a, TG13o, and TG13aR was also determined by counting colony-forming units (CFU). Viable cells were counted by gently disrupting clots and curd and spreading 10-fold serial dilutions in PBS (100 μl) on Tryptone Soy Agar (TSA) plates in technical duplicates. After incubation at 37 °C for 24 h, the CFU were determined using an automated colony counter (SphereFlash, IUL Instruments, Barcelona, Spain). The time of curd formation was determined visually as the time when curd first appeared at the bottom of wells. For some experiments, the pH was measured in the SSM media using a micro pH electrode (Thermo Scientific) every hour until the end of each experiment (up to 36 h).

### Calculation of the parameters adaptation time *τ* and maximum (absolute) growth rate *ρ_max_*

Measuring bacterial density and growth patterns using optical density (OD) is fast and reliable. However, the medium may in itself absorb light and, since the amount of absorbed light differs between media, the absolute bacterial density cannot measured. Values are instead normalised by subtracting the background OD, so that each measurement series starts at approximately zero. At later time points, equating the normalised OD to the bacterial density may be justified since the deviations from zero at the start become negligible, but that does not apply to the initial period of slow growth. Hence, only the absolute growth rate ΔX/Δt, where X is the bacterial concentration and t the time, can be calculated from the data as ΔOD/Δt for the entire series, rather than the relative or specific growth rate (ΔX/X)/Δt. (Note that using derivatives instead of differences between subsequent measurements for these calculations requires fitting of differentiable curves to the data. As the shape of our growth curves varies widely, such fitting would require further, perhaps questionable, assumptions.) We therefore measure bacterial densities at OD_600_ and calculate the parameters “adaptation time” *τ* and “maximum growth rate” *ρ_max_* from OD_600_ time series data as follows (Fig. 1B): From the empirical curve (red) differences of OD_600_ between neighbouring time points are calculated; the maximum difference corresponds to the maximum growth rate *ρ_max_* (in units of ΔOD_600_/h). The line thus defined is extended to the intersection with the x-axis. The length of time from the start of the experiment to this intersection is taken as the adaptation time *τ* (in units of h).

Furthermore, we note that measurements of bacterial concentrations via OD are only linear for low bacterial concentrations. In milk, an additional problem is clotting and curd formation at higher bacterial concentrations. We therefore only consider and show bacterial concentrations in SSM up to OD_600_ = 1. In fresh milk with 3.5% fat content, the problem is exacerbated: while the growth characteristics of TG13a (SigB-wt), TG13o and TG13aR (both SigB-def) are initially similar to those in SSM, increased clotting and curd formation compromise the measurements from about OD_600_ = 0.1, i.e. after about 4 h in TG13o and TG13aR and 10 h in TG13a (Supplementary Fig. S1E).

### Visualisation and determination of carotenoid pigmentation

The carotenoid pigmentation of *S. aureus* was visualised and the absorbance of extracts of carotenoid pigmentation measured as in a recently published article Walzl, et al. (*24*). For visualisation, a pellet that formed at the bottom of the tube after centrifugation at 10,000 *x* g for 1 min using an Eppendorf (Hamburg, Germany) centrifuge of one millilitre of overnight culture in TSB was used. The absorbance of *S. aureus* carotenoid pigmentation, including staphyloxanthin and intermediate carotenoids, was measured using a modified protocol based on methanol extraction (*75*). Briefly, overnight cultures (18 h) were grown in TSB (37 °C, shaken). Cell mass was normalised to OD_600_ = 1, and bacterial pellets were then resuspended in methanol (Carl Roth, Karlsruhe, Germany) and incubated at 55 °C for 3 min to extract pigments. The absorbance of extracted pigments was measured using a SpectraMax M3 (MolecularDevices, San Jose, CA, USA) at 300 to 750 nm intervals in 1 nm steps. Carotenoid pigmentation was determined by calculating the area under the curve (AUC) from 390 to 520 nm, with baseline adjustment using OD values at these wavelengths (OriginPro 2023, OriginLab, Northampton, MA, USA). Mean relative AUC and standard deviations were calculated from at least two independently grown samples and their technical duplicates.

### *Asp23*-expression (RT-qPCR)

Isolates were tested for *asp23*-expression (RT-qPCR). Unless stated otherwise, bacterial strains were grown in TSB (120 rpm and under aerobic conditions) and harvested at OD_600_ = 3, and *asp23*-expression was determined at the following time points: In inoculum (TSB, 18 - 20 h) for TG13a, TG13o, and TG13aR; in SSM after 2 and 4 hs for TG13a; in SSM after 8 h for TG13a, TG13o and TG13aR; and in SSM after 16, 20 and 24 h for TG13a. Depending on the actual bacterial density, batch cultures with either 15 or 50 ml of SSM were utilised to obtain adequate quantities of bacteria. Total RNA extraction from bacterial strains and *asp23*-RTqPCR was conducted following Marbach, et al. (*22*) with specific primers for *asp23* (*11*) and normalised using three reference genes (*rpoD*, *rho* and *dnaN*). Relative expression levels were determined via the REST method (*76*). Mean relative expression and standard deviations were calculated from three independently grown samples and their technical duplicates.

### Competition experiment TG13o vs TG13a

We test if the *in vivo* evolved SigB-def strain (TG13o) has a growth advantage over the initially dominant SigB-wt strain (TG13a) after transfer from TSB to SSM by co-cultivating the strains from different starting proportions of the inoculum (0.1, 0.5, and 0.9 of TG13o). Concurrently, a control experiment involving transfer of the same inocula to fresh TSB is performed. After 24, 48, 72 and 96 h, yellow and white CFUs are counted to determine the proportion of TG13o. In total, four independent iterations of this experiment are conducted, two for early (24 and 48 h) and two for late time points (72 and 96 h).

### TEM analysis

For transmission electron microscopy (TEM) examination, the SSM including the bacterial cells of TG13a (SigB-wt) and TG13o (SigB-def) and the SSM without bacteria (as control) were fixed in 5% glutaraldehyde (Merck, Darmstadt, Germany) in 0.1 M phosphate buffer (Sigma-Aldrich, St. Louis, MO, USA) at pH = 7.2 and 4 °C for 3 h. Afterwards, samples were post-fixed in 1% osmium tetroxide (Merck) in the same buffer at 4 °C for 2 h. After dehydration in an alcohol gradient series and propylene oxide (Merck), the tissue samples were embedded in glycid ether 100 (Serva, Heidelberg, Germany). Ultrathin sections were cut on a Leica Ultramicrotome (Leica Ultracut S, Vienna, Austria), stained with uranyl acetate (Sigma-Aldrich) and lead citrate (Merck), and examined on a Zeiss TEM 900 electron microscope (Carl Zeiss, Oberkochen, Germany) operated at 50 kV.

### Proteolytic activity and protein degradation

To determine the proteolytic activity in the culture media of TG13a (SigB-wt) and TG13o/TG13aR (SigB-def) in SSM, azocasein (A2765, Sigma-Aldrich), i.e., casein linked to an azo-dye, was used as a substrate for proteolytic enzymes. The breakdown of casein releases dye that we used for quantitative analysis. 250 µl of bacterial culture media was mixed with 750 µl of prewarmed azocasein working solution at 37°C. Empty SSM media without bacteria was used as a negative control. After incubation for 1 h at 37°C, the reaction was stopped by adding 100% TCA. Following incubation at room temperature for 30 min, the sample was centrifuged at 14,000 x *g* for 10 min. Absorbance at 345 nm was measured in a multi-plate reader (SpectraMax M3, MolecularDevices). Three technical duplicates of three independent experiments were analysed.

Sodium dodecyl sulfate-polyacrylamide gel electrophoresis (SDS-PAGE) and silver staining were performed to visualise the SSM degradation pattern induced by TG13a (SigB-wt) and TG13o/TG13aR (SigB-def). The culture medium was centrifuged at 19.800 x *g* for 30 min at 4°C, and proteins were concentrated using Amicon centrifugal filters with a cut-off at 100 kDa (Merck, Darmstadt, Germany) followed by Trichloroacetic acid (TCA) (Sigma-Aldrich) precipitation. The pellet was dissolved in lysis buffer (7 M urea, 2 M thiourea, 4% (w/v) CHAPS, 30 mM Tris-HCl pH 8.5), and proteins were reduced for 10 min on ice (0.5M DTT, Sigma-Aldrich), and alkylated with 14 % (w/v) iodoacetamide (Sigma-Aldrich) for 10 min at room temperature. SDS-PAGE was performed according to (*77*) using an isocratic 15%T, 18 cm x 16 cm separation gel (SE 600 Ruby, Cytiva, Marlborough, MA, USA). Protein concentration was measured by the 2D Quant kit (Cytiva). Molecular weight SDS-marker PageRuler Plus Prestained (Thermo Fisher Scientific, Waltham, MA, USA) was used as M_r_ standard, and the gel was visualised by silver staining.

### Ammonia assay

The level of ammonia/ammonium ion was determined in the culture media of TG13a (SigB-wt) and TG13o/TG13aR (SigB-def) in SSM using the Ammonia Assay Kit MAK310-1KT (Sigma-Aldrich) according to the manufacturer’s instructions. Briefly, a 100 µl sample was centrifuged for 5 min at 14,000 x *g,* and the culture media was transferred to a Vivaspin500 centrifugal concentrator (cut-off five kDa), followed by centrifugation for 45 min at 12,000 x *g*. 90 µl of the working solution was added to each 10µl flowthrough and incubated in darkness at room temperature for 15 min. The fluorescence signal was measured at 360 nm/ex and 450 nm/em in a multi-plate reader (SpectraMax M3, MolecularDevices). Technical duplicates of three independent experiments were analysed.

### Lactic acid and acetate assay

Lactic acid and acetate were determined enzymatically in the culture media of TG13a (SigB-wt) and TG13o/TG13aR (SigB-def) strains in SSM using the lactate assay kit K-LATE (Megazyme) and acetate assay kit K-ACETRM (Megazyme) according to the manufacturer’s instructions. We used the Carrez clarification set K-CARREZ (Megazyme) to remove potentially interfering substances from the SSM sample. The amount of lactic acid and acetate was determined by an enzymatic reaction monitored at 340 nm in a multi-plate reader (SpectraMax M3, MolecularDevices).

### Statistical analysis

Analyses and graphical representations were conducted in either the GraphPad Prism 10 software (10.3.1) or the statistical programming language R (*78*). Sample sizes of independent experiments are denoted as n, and data from independent experiments are always plotted. The values in the graphs for the growth curves are the mean growth at each time point plus/minus the standard deviation. When target variables were log- or logit-transformed to fulfil the assumptions of linear models this is explicitly mentioned.

To compare mean growth values across more than two groups, we generally employed an analysis of variance (ANOVA) followed by correction for multiple comparisons (Tukey’s honest significant difference or Dunnett test, as specified in each case). The competition experiment was analysed using a linear model: the logit-transformed proportion of TG13o was the target variable, and the medium (TSB vs SSM), the logit-transformed initial proportion of TG13o, and the time point were the regressors. To account for sub-structuring of variability into clonal complexes and strains within clonal complexes, mixed models with strain and clonal complex (if appropriate) as additional regressors were performed. Significance was indicated by between one and four stars when the *p*-value was below 0.05, 0.01, 0.001, and 0.0001, respectively. Details of the statistical analyses are provided in the figure legends and the main text where they occur. Some human strains hardly grew in SSM (at least during the study period), so their adaptation time and maximal growth rate could not be calculated reliably. Leaving these strains out of statistical analyses generally decreased differences between human and bovine strains and lowered the sample size, thus resulting in conservative tests. Therefore, we opted for this approach. Note that the cases where some strains did not grow at all are mentioned in the main text or figures wherever they occurred.

## Acknowledgments

We thank Stefanie Strobl, Daniela Drin, Tatjana Svoboda, Anna Walzl, Patricia Quant, Natasa Simic and Julia Schneider from the University of Veterinary Medicine Vienna for their skilful technical assistance. Fig. 1A, 1 B, and 8 were created using BioRender.com. At the time of submission TG owns a full licence to publish figures created using BioRender.

## Funding

This work was supported by

Austrian Science Fund FWF-P29304-B22 (T.G.)

National Institute of Allergy and Infectious Diseases AI124458 (L.N.S.)

National Institute of Allergy and Infectious Diseases AI157506 (L.N.S.)

## Author contributions

Conceptualisation: T.G., Si.H., C.V., L.C.M.; Data curation: T.G., C.V., L.C.M.; Formal analysis: C.V., L.C.M., T.G.; Funding acquisition: T.G., L.N.S.; Investigation: H.M., K.M-W., Se.H., L.B., A.C.P., M.-E.J., A.C.R., N.D.; Methodology: T.G., C.V., L.C.M., Si.H., L.N.S., A.C.P., R.F., W.R., N.D., I.L.; Project administration: T.G.; Resources: T.G., M.E.-S, O.K., Si.H., R.F., L.N.S., W.R.; Supervision: T.G.; Validation: C.V., L.C.M., T.G.; Visualisation: T.G., C.V., L.C.M.; Writing – original draft: T.G., C.V., L.C.M., Si.H.; Writing – review & editing: C.V., L.C.M., T.G., Si.H., M.E.-S, R.F., A.C.P., O.K.

## Data and materials availability

All datasets generated and/or analysed for this article are available from the corresponding author.

## Supplementary Materials

## Supplementary Figures

**Fig. S1.** Comparison of TG13a (SigB-wt), and TG13o and TG13aR (both SigB-def)

**Fig. S2.** In-host evolution towards SigB-deficiency in bovine mastitis isolates of various clonal complexes

**Fig. S3.** Genetic relatedness of bovine-associated CC97 and CC151 isolates

## Supplementary Tables

**Table S1.** Bovine-associated isolates from persistent IMI and engineered mutants

**Table S2.** Logistic regression analysis of the competition experiment of TG13o vs TG13a

**Table S3.** Average difference between TG13a, TG13o, and TG13aR of lactose consumption, production of metabolites, pH, and proteolytic activity after 8 h and 24 h of growth in SSM (*p*-values of t-tests shown)

**Table S4.** Bovine isolates for comparison of CC97 and CC151

**Table S5.** Human-associated isolates (CC8) and engineered mutants

**Table S6.** *S. aureus* isolates for comparison of human- and bovine-associated CC

**Table S7.** *S. aureus* Nebraska Transposon Mutant Library (NTML)

**Table S8.** The oligonucleotides employed for genetic manipulation in this research

