## Supplementary Materials for "A single transcriptional regulator is crucial for the adaptation of *Staphylococcus aureus* to diverse niches"

Claus Vogl *et al.*

**This PDF file includes:**

**Figs. S1 to S3**
**Fig. S1.** Comparison of TG13a (SigB-wt), and TG13o and TG13aR (both SigB-def)
**Fig. S2.** In-host evolution towards SigB-deficiency in bovine mastitis isolates of various clonal complexes
**Fig. S3.** Genetic relatedness of bovine-associated CC97 and CC151 isolates

**Tables S1 to S8**
**Table S1.** Bovine-associated isolates from persistent IMI and engineered mutants
**Table S2.** Logistic regression analysis of the competition experiment of TG13o vs TG13a
**Table S3.** Average difference between TG13a, TG13o, and TG13aR of lactose consumption, production of metabolites, pH, and proteolytic activity after 8 h and 24 h of growth in SSM (*p*-values of t-tests shown)
**Table S4.** Bovine isolates for comparison of CC97 and CC151
**Table S5.** Human-associated isolates (CC8) and engineered mutants
**Table S6.** *S. aureus* isolates for comparison of human- and bovine-associated CC
**Table S7.** *S. aureus* Nebraska Transposon Mutant Library (NTML)
**Table S8.** The oligonucleotides employed for genetic manipulation in this research

**Fig. S1.**

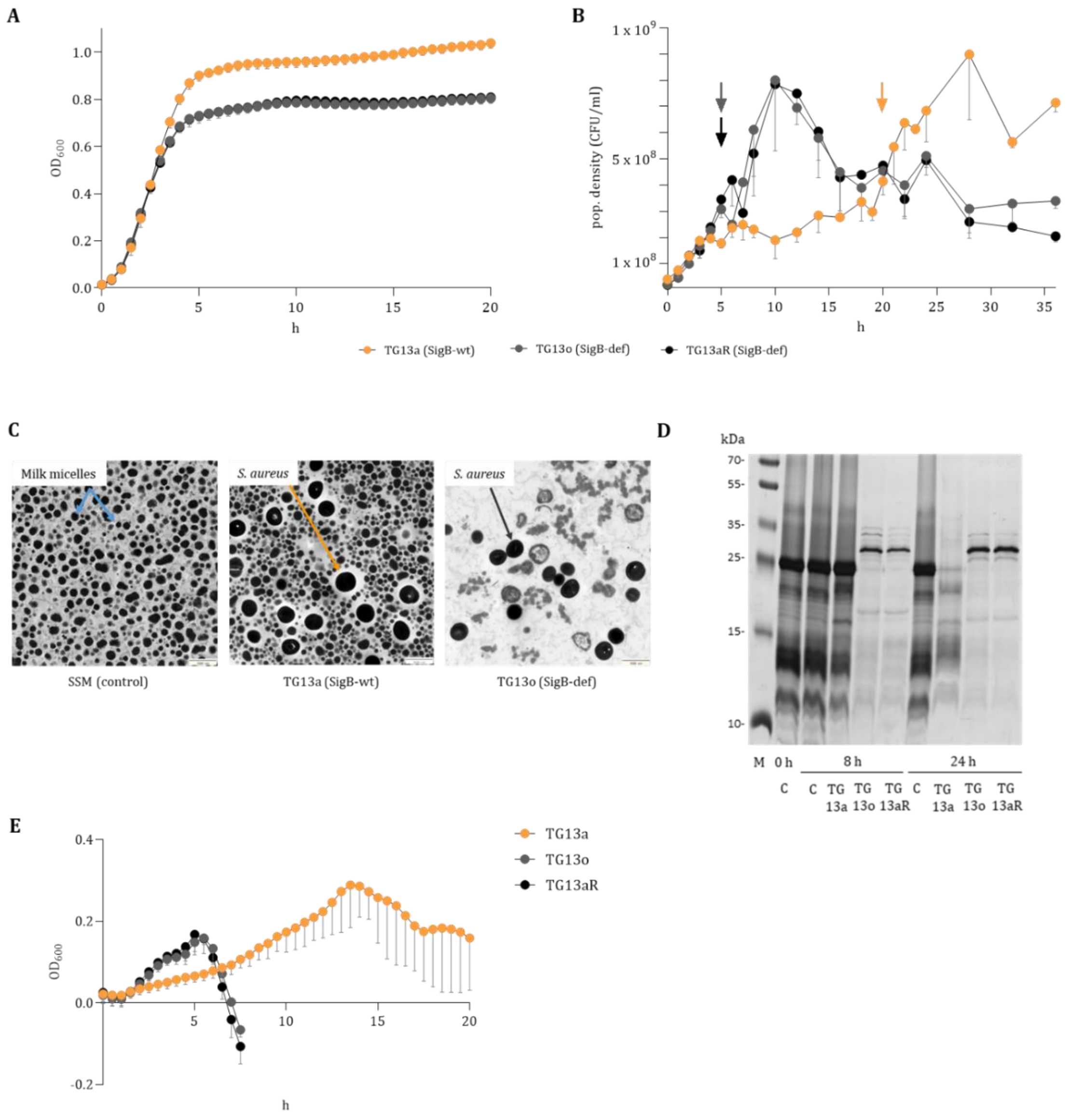


**Comparison of TG13a (SigB-wt), and TG13o and TG13aR (both SigB-def). (A)** *S. aureus* strains pre-cultivated in TSB and then transferred to fresh TSB grow without an adaptation phase. **(B)** Population density in CFU/ml after transfer to SSM. The arrows mark the time of curd formation. **(C)** TEM images of SSM without bacteria (media control), and after growth for 24 h in SSM. **(D)** SDS-PAGE (15%) of the supernatant of bacteria grown in SSM and harvested at the indicated time. Media controls without bacteria, labelled C, are included at time points t = 0, t = 8 and t =24. Sample load was normalised to equal protein concentration. **(E)** The growth curves ($OD_{600}$) of TG13a (SigB-wt), TG13o, and TG13aR (both SigB-def) after transfer from TSB to fresh milk with 3.5% fat content ($OD_{600}$; mean – SD).

M, Marker proteins. TG13a, SigB-wt, TG13o and TG13aR, both SigB-def.

**Fig. S2.**

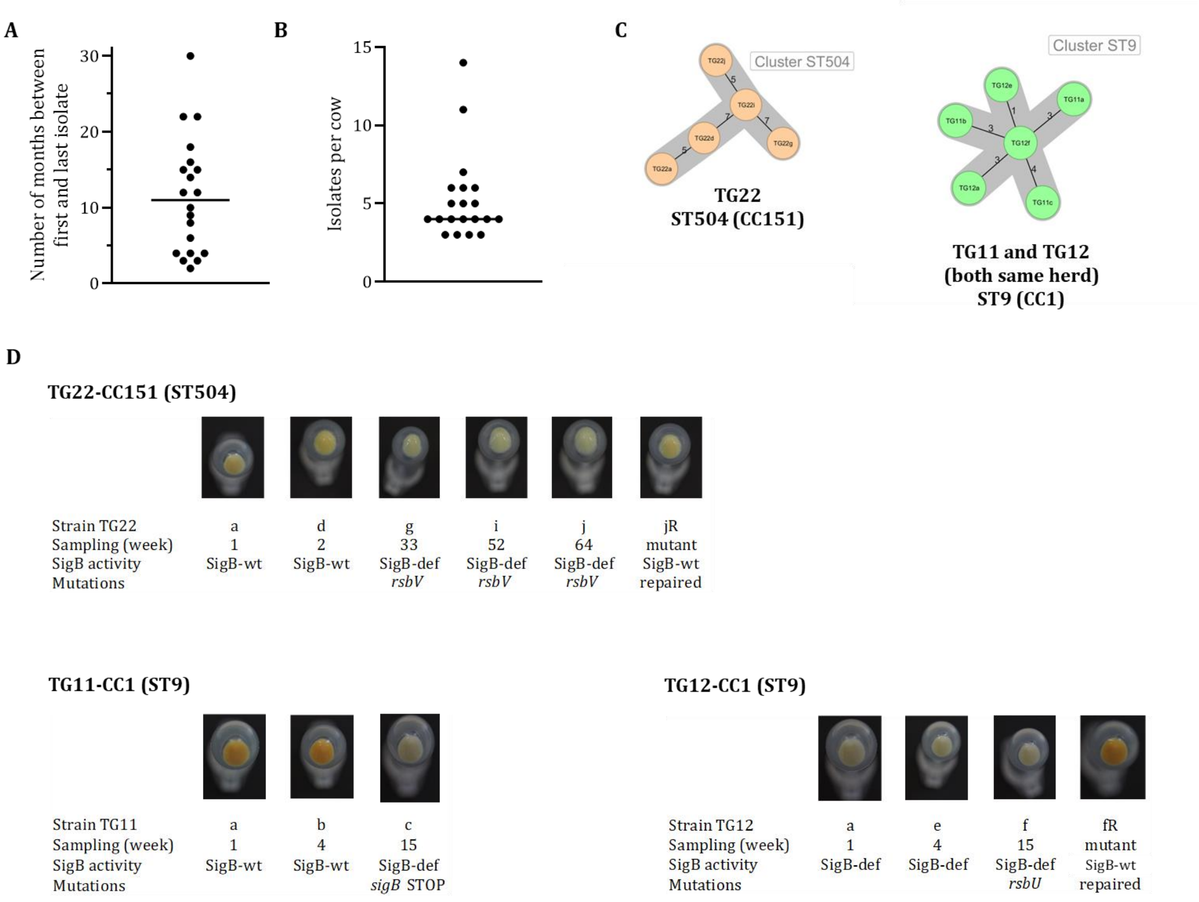


**In-host evolution towards SigB-deficiency in bovine mastitis isolates of various clonal complexes. (A)** Intervals between the first and the last isolate collected from individual dairy cows (n = 20), and **(B)** number of isolates collected from each cow (from which a minimum of three isolates was obtained) by retrospective screening for SigB-def identifiable by visible loss of colony pigmentation (i.e., for a white or grey instead of a yellow colony colour) from consecutively sampled isolates of *S. aureus* from bovine hosts with persistent IMI (n = 20). **(C)** Putative SigB-def isolates in milk samples from three cows from two herds. Minimum spanning tree using core genome multilocus sequence typing (cgMLST) data for TG22 and TG11/TG12 (same herd) suggests clonality, although the isolates were recovered from different cows, suggesting transmission from one cow to the other. We used a cluster threshold of ≤24 allelic differences in the core genome to determine clonality. For TG22, the first and last isolates were collected 15 months apart. Both of these, as well as the three intermediate isolates collected from the same cow, have a cgMLST cluster of less than 24 allelic differences, indicating clonality. Isolates were assigned to the clonal complexes CC97, CC151, and CC1. **(D)** SigB activity: In both reconstructed strains (TG22jR and TG12jR) carotenoid pigmentation is restored, confirming that the mutations are solely responsible for the inactivation of the *sigB* operon.

**Fig. S3.**

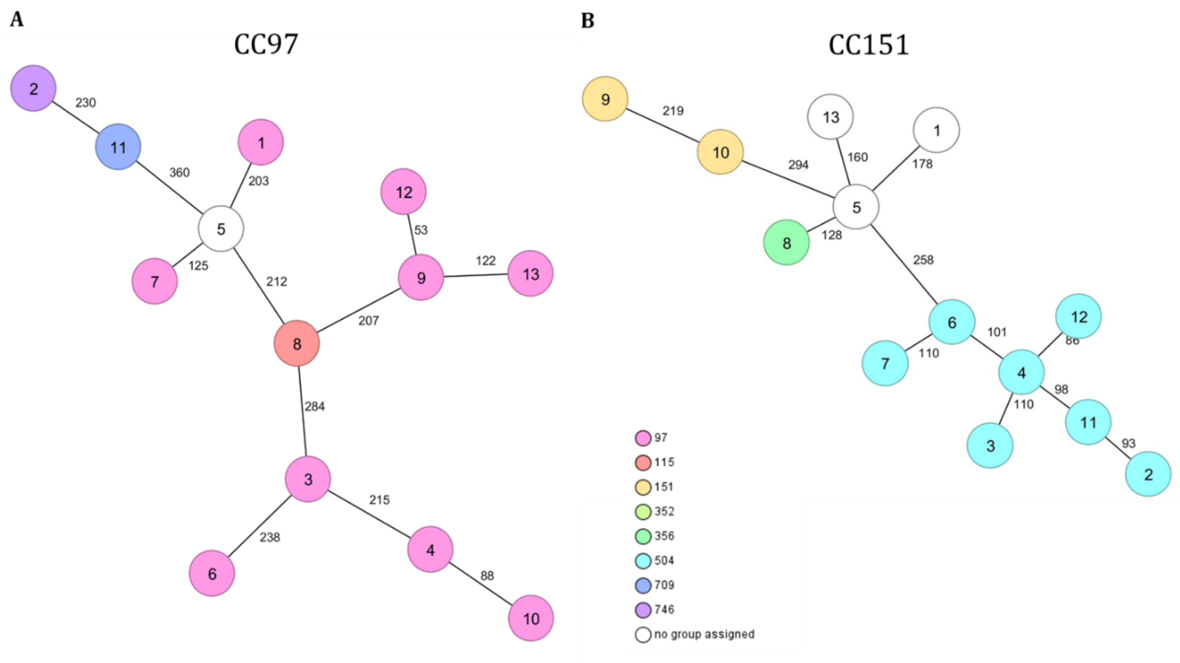


**Genetic relatedness of bovine-associated CC97 and CC151 isolates.** Minimum spanning tree using core genome multilocus sequence typing (cgMLST) data for strains of **(A)** CC97 (n = 13) and **(B)** CC151 (n = 13) isolated from the milk of cows from different geographic regions (see the list of strains and their characteristics in Supplementary Table 4). Graphical representation of their relationships obtained by a cgMLST analysis. All isolates show unique cgMLST patterns, exceeding the threshold of ≤24 allelic differences in the core genome used to determine clonality. The colour of the nodes indicates the ST type.

**Table S1.**

**Bovine-associated isolates from persistent IMI and engineered mutants**

| **Strain** | **Description** | **SigB activity[1]** | **Sampling week** | **CC** | **ST** | ***spa*-type** | ***cap-gene*** | ***agr*-type** | **Reference** |
|---|---|---|---|---|---|---|---|---|---|
| TG13a | First isolate (IN) | + | 1 | | | | | | Marbach *et al.* 2019 |
| TG13aΔ*sspA* | *sspA* mutant of TG13a | + | | | | | | | this study |
| TG13aΔ*aur* | *aur* mutant of TG13a | + | | | | | | | this study |
| TG13o | 21$^{th}$ and last isolate (HA), incl. mutation in *rsbU* (G368A) | - | 14 | | | | | | Marbach *et al.* 2019 |
| TG13oΔ*sspA* | *sspA* mutant of TG13o | - | | 97 | 352 | t044 | 5 | *I* | this study |
| TG13oΔ*aur* | *aur* mutant of TG13o | - | | | | | | | this study |
| TG13aR | TG13a::*rsbU* (G368A) mutant | - | | | | | | | Walzl *et al.* 2023 |
| TG13aRΔ*sspA* | *sspA* mutant of TG13aR | - | | | | | | | this study |
| TG13aRΔ*aur* | *aur* mutant of TG13aR | - | | | | | | | this study |
| **TG11-CC1** | | | | | | | | | |
| TG11a | First isolate | + | 1 | | | | | | this study |
| TG11b | Second isolate | + | 4 | 1 | 9 | t3446 | 5 | *II* | this study |
| TG11c | Third and last isolate, incl. stop codon in *sigB* | - | 15 | | | | | | this study |
| **TG12-CC1** | | | | | | | | | |
| TG12a | First isolate, incl. SNP G431T in *rsbU* | - | 1 | | | | | | this study |
| TG12e | Second isolate, incl. SNP G431T in *rsbU* | - | 4 | 1 | 9 | t3446 | 5 | *II* | this study |
| TG12f | Third and last isolate, incl. SNP G431T in *rsbU* | - | 15 | | | | | | Walzl *et al.* 2023 |
| TG12fR | TG12f::*rsbU* repaired mutant | + | | | | | | | Walzl *et al.* 2023 |
| **TG22-CC151** | | | | | | | | | |
| TG22a | First isolate | + | 1 | | | | | | Grunert *et al.* 2018 |
| TG22d | Second isolate | + | 2 | | | | | | Grunert *et al.* 2018 |
| TG22g | Third isolate, incl. 62bp trunc. in *rsbV* | - | 33 | 151 | 504 | t529 | 8 | *II* | Grunert *et al.* 2018 |
| TG22i | Fourth isolate, incl. 62bp trunc. in *rsbV* | - | 52 | | | | | | Grunert *et al.* 2018 |
| TG22j | Fifth and last isolate, incl. 62bp trunc. in *rsbV* | - | 64 | | | | | | Grunert *et al.* 2018 |
| TG22jR | TG22j::*rsbV* repaired mutant | + | | | | | | | this study |

[1)] +, SigB-wt; -, SigB-deficient

### Table S2.

**Logistic regression analysis of the competition experiment of TG13o vs TG13a**: Inncolum with different proportions of TG13o (0.1, 0.5, 0.9) was grown in both TSB and SSM, and the growth of each strain was evaluated at different time points (24h, 48h, 72h, 96h). The dependent variable in the regression analysis is the log-odds of the observed strain proportions. This is regressed onto the independent variables encoding the medium (TSB corresponds to 0; SSM to 1), the evaluation times, and the log-odds of the initial strain proportions (note this precludes inclusion of an intercept). Including a random effect term for different replicates does not improve model fit (not shown).

```
Call:
lm(formula = logit(proportion) ~ 0 + times * medium * logit(init pr),
    data = df_biol)
Residuals:
    Min      1Q  Median      3Q     Max
-1.5657 -0.4528  0.1640  0.4504  1.6048
Coefficients:
                                 Estimate Std. Error t value Pr(>|t|)
times                             0.001748   0.005398   0.324 0.747714
medium_TSB                       -0.508199   0.354815  -1.432 0.159830
medium_SSM                       -0.565521   0.354815  -1.594 0.118843
logit(init pr)                    0.915966   0.197776   4.631 3.8e-05  ***
times:medium_SSM                  0.028384   0.007634   3.718 0.000615 ***
times:logit(init pr)             -0.006171   0.003009  -2.051 0.046875 *
Medium_SSM:logit(init pr)         0.012214   0.279697   0.044 0.965385
times:medium_SSM:logit(init pr)  0.001241   0.004255   0.292 0.772111
---
Signif. codes:  0 '***' 0.001 '**' 0.01 '*' 0.05 '.' 0.1 ' ' 1
Residual standard error: 0.7096 on 40 degrees of freedom
Multiple R-squared:  0.8499,    Adjusted R-squared:  0.8199
F-statistic: 28.31 on 8 and 40 DF,  p-value: 3.759e-14
```

### Table S3.

**Average difference between TG13a, TG13o, and TG13aR of lactose consumption, production of metabolites, *pH*, and proteolytic activity after 8 h and 24 h of growth in SSM (*p*- values of t-tests shown)**

| | 8_p_overall | 8_p_13o-13a | 8_p_13rA-13a | 8_p_13rA-13o | 24_p_overall | 24_p_13o-13a | 24_p_13rA-13a | 24_p_13rA-13o |
|---|---|---|---|---|---|---|---|---|
| **pH** | 2.843171e-03 | 4.499992e-03 | 5.008966e-03 | 0.993082793 | 9.619389e-04 | 0.0018166542 | 0.0015131405 | 0.9733890 |
| **acetate** | 2.002997e-04 | 4.349719e-04 | 2.957351e-04 | 0.831265171 | 5.130239e-04 | 0.0009346894 | 0.0008547612 | 0.9922753 |
| **lactate** | 2.521679e-05 | 4.469903e-05 | 4.614951e-05 | 0.997530486 | 7.809455e-05 | 0.0001280476 | 0.0001530213 | 0.9474261 |
| **proteolytic activity** | 1.334007e-04 | 2.334014e-04 | 2.408150e-04 | 0.998577416 | 4.161395e-03 | 0.0056157065 | 0.0087043103 | 0.7803790 |
| **ammonia** | 3.153807e-06 | 2.723864e-06 | 1.952173e-05 | 0.004176723 | 1.009559e-04 | 0.0003202220 | 0.0001193526 | 0.2958623 |

### Table S4.

#### Bovine isolates for comparison of CC97 and CC151

| CC | ID | ST | *spa-* type | *cap -* gene | *agr-* type | Strain | Origin[1] | Reference |
|---|---|---|---|---|---|---|---|---|
| | 1 | ST97 | t267 | | | A1 | ARG | Grunert *et al.* 2013 |
| | 2 | ST746 | t267 | | | A4 | ARG | Grunert *et al.* 2013 |
| | 3 | ST97 | t2112 | | | 39FR | RWA | Antok *et al.* 2019 |
| | 4 | ST97 | t1236 | | | 87RL | RWA | Antok *et al.* 2019 |
| | 5 | ST97 | t521 | | | MB328 | ARG | Tuchscherr *et al.* 2007 |
| | 6 | ST97 | t10103 | | | 101RR | RWA | Antok *et al.* 2019 |
| CC97 | 7 | ST97 | t521 | 5 | *I* | A10 | ARG | Grunert *et al.* 2013 |
| | 8 | ST115 | t044 | | | N305 | CAN | Prasad *et al.* 1968 |
| | 9 | ST97 | t044 | | | S413-1 | AUT | Kümmel *et al.* 2016 |
| | 10 | ST97 | t9432 | | | 37RR | RWA | Antok *et al.* 2019 |
| | 11 | ST709 | t267 | | | MB325 | ARG | Tuchscherr *et al.* 2007 |
| | 12 | ST97 | t044 | | | S387-2 | AUT | Kümmel *et al.* 2016 |
| | 13 | ST97 | t267 | | | 127 | AUT | Schabauer *et al.* 2018 |
| | 1 | ST705 | t529 | | | A20 | ARG | Grunert *et al.* 2013 |
| | 2 | ST504 | t529 | | | 20 | AUT | Schabauer *et al.* 2018 |
| | 3 | ST504 | t529 | | | 14c | AUT | this study |
| | 4 | ST504 | t529 | | | 38 | AUT | Schabauer *et al.* 2018 |
| | 5 | ST705 | t529 | | | A18 | ARG | Grunert *et al.* 2013 |
| | 6 | ST504 | t529 | | | S392-1 | AUT | Kümmel *et al.* 2016 |
| CC151 | 7 | ST504 | t529 | 8 | *II* | S429 | AUT | Kümmel *et al.* 2016 |
| | 8 | ST356 | t529 | | | A14 | ARG | Grunert *et al.* 2013 |
| | 9 | ST151 | t529 | | | 115 | AUT | Schabauer *et al.* 2018 |
| | 10 | ST151 | t529 | | | RF122 | GBR | Fitzgerald *et al.* 2000 |
| | 11 | ST504 | t529 | | | LMM-22 | AUT | this study |
| | 12 | ST504 | t529 | | | W18c | AUT | Grunert *et al.* 2018 |
| | 13 | ST705 | t458 | | | A15 | ARG | Grunert *et al.* 2013 |

1) IRL, Irland; ARG, Argentina; RWA, Rwanda; CAN, Canada; AUT; Austria; GBR, United Kingdom.

| | | | | | | | | |
|---|---|---|---|---|---|---|---|---|
| CC151 | MOK124-CC151 | ST151 | nd | 8 | *II* | MOK124 | IRL | Murphy *et al.* 2019 |
| CC97 | MOK023-CC97 | ST3170 | nd | 5 | *I* | MOK023 | IRL | Murphy *et al.* 2019 |

**Table S5.**

**Human-associated isolates (CC8) and engineered mutants**

| Strain | Description | SigB activity[1] | CC | ST | *spa* -type | *cap-gene* | *agr* -type | Reference |
|---|---|---|---|---|---|---|---|---|
| Nm | Newman - human infection | + | 8 | 254 | t008 | 5 | nd | Duthi *et al.* 1952 |
| NmR | Newman::*rsbU* (G368A) mutant | - | | | | | | Walzl *et al.* 2023 |
| 8325-4 | 11 bp deletion in *rsbU* gene | - | | | | | | Novick *et al.* 1967 |
| SH1000 | *rsbU* repaired mutant | + | 8 | 8 | t211 | 5 | nd | Horsburgh *et al.* 2002 |
| SH1000Δ*sigB* | *sigB* deletion mutant of SH1000 | - | | | | | | Tuchscherr *et al.* 2015 |
| LAC | USA300 LAC (AH1263) | + | | | | | | Boles *et al.* 2010 |
| LACΔ*sigB* | USA300 LAC *sigB* ::tet (LNS2602) | - | 8 | 8 | nd | 5 | nd | this study |
| ESPNΔ*sigB* | USA300 LAC Δ*aur* Δ*sspAB* Δ*scpA* spl::erm *sigB* ::tet (LNS4021) | - | | | | | | this study |

1) +: SigB-wt; -: SigB-deficient

**Table S6.**

***S. aureus* isolates for comparison of human- and bovine-associated CC**

| Host | CC | ST | *spa*-type | *cap*-gene | *agr*-type | Strain | Reference |
|---|---|---|---|---|---|---|---|
| HUMAN | CC8 | ST250 | t008 | 5 | *I* | Newman | Duthi *et al.* 1952 |
| | CC8 | ST254 | t008 | 5 | *I* | COL | Dyke *et al.* 1966 |
| | CC8 | ST8 | t648 | 5 | *I* | 52 | Johler *et al.* 2016 |
| | CC8 | ST72 | t148 | 5 | *I* | 29 | Johler *et al.* 2016 |
| | CC30 | nd | t018 | 5 | *III* | 16 | Johler *et al.* 2016 |
| | CC30 | ST30 | t021 | 5 | *III* | 43 | Johler *et al.* 2016 |
| | CC30 | ST36 | t018 | 5 | *III* | 37 | Johler *et al.* 2016 |
| | CC5 | ST4255 | t8456 | 5 | *II* | 39 | Johler *et al.* 2016 |
| | CC5 | ST5 | t857 | 5 | *II* | 57 | Johler *et al.* 2016 |
| | CC5 | ST5 | t179 | 5 | *II* | 38 | Johler *et al.* 2016 |
| | CC45 | ST508 | t630 | 8 | *I* | 19 | Johler *et al.* 2016 |
| | CC45 | ST45 | t950 | 8 | *I* | 27 | Johler *et al.* 2016 |
| | CC45 | ST45 | t445 | 8 | *I* | 68 | Johler *et al.* 2016 |
| BOVINE | CC151 | ST504 | t529 | 8 | *II* | S392-1 | Kümmel *et al.* 2016 |
| | CC151 | ST151 | t529 | 8 | *II* | RF122 | Fitzgerald *et al.* 2000 |
| | CC151 | ST504 | t529 | 8 | *II* | 14c | this study |
| | CC97 | ST115 | t044 | 5 | *I* | N305 | Prasad *et al.* 1968 |
| | CC97 | ST709 | t267 | 5 | *I* | MB325 | Tuchscherr *et al.* 2007 |
| | CC97 | ST97 | t044 | 5 | *I* | S387-2 | Kümmel *et al.* 2016 |
| | CC8 | ST8 | t2953 | 5 | *I* | S276-2 | Kümmel *et al.* 2016 |
| | CC8 | ST8 | t2953 | 5 | *I* | LMM-216 | this study |
| | CC8 | ST8 | t2953 | 5 | *I* | 108 | Schabauer *et al.* 2018 |
| | CC8 | ST8 | t2953 | 5 | *I* | 171 | Schabauer *et al.* 2018 |

### Table S7.

#### *S. aureus* Nebraska Transposon Mutant Library (NTML)

NTML mutants are based on human-derived, wild-type JE2, a plasmid-cured variant of USA300 LAC (CC8)

| # | Gene ID SAUSA300_ | gene name | proteolytic capacity ($log_2$ fold change compared to WT) after Giemza *et al.* 2019 |
|---|---|---|---|
| 1 | 0063 | 0063 | -0,2630 |
| 2 | 0066 | argR1 | -1,7655 |
| 3 | 0093 | 0093 | -0,5850 |
| 4 | 0095 | 0095 | 0,2630 |
| 5 | 0104 | 0104 | 0,7655 |
| 6 | 0110 | norG | 0,1375 |
| 7 | 0114 | sarS | 0,9260 |
| 8 | 0137 | 0137 | 0,0000 |
| 9 | 0195 | rpiRB | 1,1375 |
| 10 | 0217 | 0217 | -0,5850 |
| 11 | 0238 | 0238 | -0,2630 |
| 12 | 0255 | lytR | 0,1375 |
| 13 | 0258 | 0258 | 0,0000 |
| 14 | 0265 | rbsR | -0,3785 |
| 15 | 0315 | nanA | 0,2630 |
| 16 | 0333 | 0333 | -0,1375 |
| 17 | 0334 | mepR | -0,2630 |
| 18 | 0373 | 0373 | 0,5850 |
| 19 | 0473 | purR | -0,1375 |
| 20 | 0503 | 0503 | -0,1375 |
| 21 | 0519 | sigH | 0,4854 |
| 22 | 0577 | 0577 | 0,2630 |
| 23 | 0621 | mntR | -2,2016 |
| 24 | 0645 | graR | 0,7655 |
| 25 | 0653 | rbf | -0,7655 |
| 26 | 0654 | sarX | 0,6781 |
| 27 | 0658 | ccpE | -0,7655 |
| 28 | 0691 | saeR | -0,6781 |
| 29 | 0954 | atlR | -2,1043 |
| 30 | 0998 | 0998 | 0,6781 |
| 31 | 1019 | 1019 | -0,6781 |
| 32 | 1148 | codY | 1,3219 |
| 33 | 1170 | 1170 | 0,0000 |
| 34 | 1204 | 1204 | 0,2630 |
| 35 | 1220 | 1220 | 0,4854 |
| 36 | 1253 | 1253 | 0,0000 |
| 37 | 1308 | arlR | 0,1375 |
| 38 | 1409 | vraR | 0,2630 |
| 39 | 1424 | 1424 | -0,3785 |
| 40 | 1433 | 1433 | -0,2630 |
| 41 | 1442 | srrA | 0,1375 |
| 42 | 1448 | fur | -0,2630 |
| 43 | 1455 | 1455 | -0,9260 |
| 44 | 1457 | malR | -0,1375 |
| 45 | 1469 | argR | 0,5850 |
| 46 | 1514 | zur | -0,1375 |
| 47 | 1542 | hrcA | -0,2630 |
| 48 | 1583 | cymR | 0,2630 |
| 49 | 1639 | phoP | 0,2630 |
| 50 | 1682 | ccpA | -0,2630 |
| 51 | 1708 | rot | 0,7655 |
| 52 | 1717 | arsR | 0,2630 |
| 53 | 1722 | sigS | 0,2630 |
| 54 | 1797 | xdrA | 0,5850 |
| 55 | 1842 | perR | 0,0000 |
| 56 | 1946 | 1946 | -0,1375 |
| 57 | 1968 | 1968 | -0,2630 |
| 58 | 1969 | 1969 | -0,8480 |
| 59 | 1992 | agrA | -2,5607 |
| 60 | 1995 | scrR | -0,3785 |
| 61 | 1999 | rex | -0,9260 |
| 62 | 2022 | sigB | 1,3219 |
| 63 | 2036 | kdpE | 0,2630 |
| 64 | 2050 | 2050 | -0,7655 |
| 65 | 2098 | czrA | 0,0000 |
| 66 | 2106 | mtlR | -0,6781 |
| 67 | 2156 | lacR | -0,9260 |
| 68 | 2160 | 2160 | 0,0000 |
| 69 | 2216 | 2216 | -0,1375 |
| 70 | 2247 | sarY | 0,6781 |
| 71 | 2248 | 2248 | -1,5850 |
| 72 | 2264 | rpiRC | 0,3785 |
| 73 | 2271 | hex | 0,0000 |
| 74 | 2279 | hutR | -0,6781 |
| 75 | 2300 | 2300 | 0,3785 |
| 76 | 2303 | tcaR | 0,1375 |
| 77 | 2308 | hssR | 0,2630 |
| 78 | 2310 | 2310 | -0,6781 |
| 79 | 2322 | 2322 | 0,8480 |
| 80 | 2326 | rsp | -0,3785 |
| 81 | 2331 | sarZ | 0,6781 |
| 82 | 2336 | 2336 | 0,0000 |
| 83 | 2337 | nrec | 0,2630 |
| 84 | 2437 | sarT | -0,8480 |
| 85 | 2438 | sarU | -0,6781 |
| 86 | 2445 | 2445 | -1,0704 |
| 87 | 2459 | 2459 | -0,8480 |
| 88 | 2480 | cidR | -0,3785 |
| 89 | 2490 | 2490 | 0,2630 |
| 90 | 2509 | 2509 | -1,0000 |
| 91 | 2515 | 2515 | 0,0000 |
| 92 | 2530 | 2530 | 0,1375 |
| 93 | 2547 | 2547 | -0,8480 |
| 94 | 2559 | nsaR | 0,8480 |
| 95 | 2566 | arcR | -0,1375 |
| 96 | 2571 | argR3 | -0,1375 |
| 97 | 2575 | 2575 | 0,0000 |
| 98 | 2599 | icaR | 0,4854 |
| 99 | 2640 | 2640 | 0,1375 |

**The following transcriptional regulator mutants were excluded from analysis, because they did not grow in SSM within 72 h**

| | | |
|---|---|---|
| 100 | 444 | gtlC |
| 101 | 450 | treR |
| 102 | 605 | sarA |
| 103 | 1174 | 1174 |

**The following transcriptional regulator mutants were not available at time of acquisition**

| | | |
|---|---|---|
| 1 | 804 | 804 |
| 2 | 1798 | airR |
| 3 | 1888 | hisR |
| 4 | 2218 | sarV |
| 5 | 2425 | 2425 |

**Table S8.**

**The oligonucleotides employed for genetic manipulation in this research**

| Primer name | 5'-3' sequence | Description |
|---|---|---|
| OL7735 | ATGggatccGAGCAGGTGCGAAATAATGG | *sigB* confirmation F |
| OL7736 | ATGgaattcCAAATTCTATTGATGTGCTGCTTCTTG | *sigB* confirmation R |
| *aur* A | cctcactaaagggaacaaaagctgggtaccCATTAGGCATCTGGTTTGTC | *aur* deletion construct generation in pIMAY-Z |
| *aur* B | gtttaacattacttcttcttgtttatTTCCTCCTGAAATCTTAAAAACAG | *aur* deletion construct generation in pIMAY-Z |
| *aur* C | ctgtttttaagatttcaggaggaaATAAACAAGAAGAAGTAATGTTAAAC | *aur* deletion construct generation in pIMAY-Z |
| *aur* D | cgactcactatagggcgaattggagctcTATGAACCATTGATGATTGAACT | *aur* deletion construct generation in pIMAY-Z |
| *aur* OUT F | AACGCGATTAAGTATGAT | Screen for *aur* deletion mutants |
| *aur* OUT R | CCAGGTGAGGTTTTGAC | Screen for *aur* deletion mutants |
| *sspA* A | cctcactaaagggaacaaaagctgggtaccGGACGTCGTGAACTA | *sspA* deletion construct generation in pIMAY-Z |
| *sspA* B | tactaaatctaaattaagatgaagtttaCATCTAAAAACCTCCAAAAAA | *sspA* deletion construct generation in pIMAY-Z |
| *sspA* C | ttttttggaggtttttagatgTAAACTTCATCTTAATTTAGATTTAGTA | *sspA* deletion construct generation in pIMAY-Z |
| *sspA* D | attggagctccaccgcggtggcggccgcTTTAGCACTTTCTTTTTCTTTTACA | *sspA* deletion construct generation in pIMAY-Z |
| *sspA* OUT F | GCAATCGTTCCAGGCTCATC | Screen for *sspA* deletion mutants |
| *sspA* OUT R | TCTTCTTGTATCGCTTCGTTTTC | Screen for *sspA* deletion mutants |
| pIMAY-Z MCS F | TACATGTCAAGAATAAACTGCCAAAGC | Screen pIMAY-Z constructs |
| pIMAY-Z MCS R | AATACCTGTGACGGAAGATCACTTCG | Screen pIMAY-Z constructs |
| *rsbV*_Trunc_F | AGGGAACAAAAGCTGGGTACCTTAAAAAACATAAACATATGCACC | Repair 61-bp truncation in *rsbV* |
| *rsbV*_Trunc_R | ATAGGGCGAATTGGAGCTCTTATATACTTAGTCATACTGATTGTC | Repair 61-bp truncation in *rsbV* |
